# Genome organizer SATB1 restrains the BACH1-cMAF axis that drives autoimmunity

**DOI:** 10.1101/2025.11.25.690242

**Authors:** Dionysios Alexandros Papamatheakis, Petros Tzerpos, Despina Tsoukatou, Eleftherios Morres, Manοuela Kapsetaki, Dezso Balázs, Kazuhiko Igarashi, Charalampos Spilianakis

**Author notes:** Senior author (Charalampos Spilianakis).

## Abstract

The establishment of central tolerance in the thymus is governed by precise gene regulatory networks, but how disrupted chromatin architecture leads to autoimmunity is unclear. The genome organizer SATB1 is essential for T cell development and its loss triggers a severe autoimmune phenotype. Here, we identify a pathogenic transcriptional axis, involving BACH1 and cMAF, that is unleashed upon SATB1 deletion. Using integrated multi-omics in T cell-specific *Satb1*-knockout mice, we demonstrate that SATB1 constrains BACH1 chromatin occupancy. In its absence, BACH1 redistributes to promoter-proximal regions and SATB1-bound immune loci, where it facilitates the recruitment of the transcription factor cMAF. This BACH1-cMAF complex drives a pro-inflammatory transcriptional program in thymocytes, which seeds the periphery and results in a systemic autoimmune disease. Strikingly, genetic ablation of *Bach1* in *Satb1*-deficient mice rescues the pathology, normalizes immunity and prevents mortality. Furthermore, genetic or pharmacological inhibition of cMAF ameliorates the disease. The pro-inflammatory signature in mutant T cells overlaps with T cells from Systemic Lupus Erythematosus (SLE) patients. Our findings reveal a BACH1-cMAF axis that is derepressed upon SATB1 loss and bridges disrupted thymic chromatin organization to peripheral autoimmunity, nominating new therapeutic targets.

## Introduction

The immune system must constantly balance responsiveness to pathogens with tolerance to self. Failure to maintain this equilibrium allows autoreactive lymphocytes to persist and drive chronic inflammation, culminating in autoimmune diseases such as systemic lupus erythematosus (SLE), multiple sclerosis and rheumatoid arthritis. These disorders share common features of dysregulated T cell development, aberrant cytokine production and sustained activation of proinflammatory pathways. Although genome-wide association studies have identified risk loci and effector pathways in human autoimmunity, how altered transcriptional control, during thymic development, translates into pathogenic T cell states in the periphery remains unclear.

This equilibrium is governed by the proper expression of T-cell specific surface markers (i.e. CD3, CD4 and CD8), signaling molecules (i.e. LCK, ZAP70 and ITK) and lineage specification transcription factors (i.e. TCF-1, BCL11b and GATA3)^1–4^. Additionally, all developing T cells need to express properly recombined T-cell receptors (TCRs) in order to identify a wide variety of exogenous antigens^5^. The aforementioned features that characterize proper T cell development and function are governed by the correct spatiotemporal gene expression during commitment and differentiation of T cells in the thymus^6,7^. The different developmental stages during thymic development are governed by the combinatorial role of tissue specific transcription factors and cell-cell communication in the thymus^8,9^.

The chromatin organizer Special AT-rich sequence-binding protein 1 (SATB1) exemplifies a transcriptional regulator essential for T cell development. SATB1 binds base-unpairing or matrix attachment regions and recruits chromatin remodelers to establish higher-order loops. Recent studies have highlighted that SATB1 safeguards T cell development and its loss leads to the deregulation of T cell development leading to ectopic T cell activation and eventually to an autoimmune-like phenotype^10,11^. It is highly expressed at the double positive CD4^+^CD8^+^ (DP) stage of thymocyte maturation, where it contributes to lineage choice and negative selection^12^. Loss of SATB1 specifically in the T cell compartment results in a striking autoimmune-like phenotype: CD4⁺ and CD8⁺ T cells become dysregulated, infiltrate multiple tissues, produce autoantibodies and lead to premature mortality^10^. Collectively, SATB1 has been established as an essential gatekeeper of chromatin topology and tolerance in the thymus.

Using DNA-affinity purification assays coupled to mass spectrometry analysis, we previously found that a biotinylated probe corresponding to the *Rad50* DNase I hypersensitive site 6 (RHS6) region of the T helper type 2 (Th2) locus specifically captured both SATB1 and BTB and CNC homology factor 1 (BACH1), indicating that these proteins strongly associate with this genomic element^13^. BACH1 is a member of the CNC-bZIP transcription factor family, broadly expressed in multiple tissues and was initially characterized as a heme-responsive repressor. It is a crucial regulator of oxidative stress response, identified as the repressor of the *Hmox1* gene in normoxia^14^. The role of BACH1 in T cell biology still remains obscure. BACH1 plays an important role in macrophage-mediated osteoclastogenesis leading to rheumatoid arthritis and its loss leads to decreased osteoclast destruction, reduced *Tnf*α expression and reduced inflammatory bone loss through derepression of *Hmox1* and *Blimp1*^15,16^. *Bach1*^−/-^ mice, in which arthritis has been induced, have decreased joint lesions compared to their wild type counterparts upon experimentally induced inflammation. Specifically, loss of BACH1 leads to M2 macrophage polarization that facilitate the downregulation of proinflammatory cytokine production and reduction of neutrophil infiltration in mice with experimental colitis^17^.

BACH1 cooperates with small MAFs (MAFF, MAFK), at MAF-recognition sites (MAREs) to regulate gene expression^18–20^. cMAF belongs to the AP-1/bZIP family and is a versatile regulator of T cell biology. It has been studied primarily in peripheral effector and regulatory T cell subsets. In Th17 and T follicular helper cells, cMAF promotes IL-21 production and supports effector differentiation^21,22^; in regulatory T cells and Tr1 cells^23^, it drives IL-10 expression^24^, conferring immunoregulatory capacity^25–28^. cMAF therefore straddles pathogenic and regulatory functions depending on cellular context. However, the upstream cues that determine where cMAF binds in the genome remain incompletely understood and whether cMAF contributes to autoimmune pathogenesis from the earliest stages of T cell development has not been explored.

Taken together, SATB1, BACH1 and cMAF highlight three dimensions of transcriptional control: chromatin organization, stress-sensing transcriptional regulation and effector cytokine regulation. Yet their intersections remain uncharted. SATB1 and BACH1 have both been described as transcriptional regulators, but whether their functions are cooperative, competitive, or insulated from one another is unknown. While genetic risk loci and effector pathways have been described in human autoimmune syndromes, the molecular events that connect altered transcriptional control during lymphocyte development to pathogenic effector functions in the periphery remain poorly defined.

In this study, we address this gap by integrating chromatin and transcriptomic profiling of thymocytes and peripheral lymphocytes across *Satb1*- and *Bach1*-deficient mouse models. We show that SATB1 normally restrains BACH1 occupancy at immune-regulatory loci. When SATB1 is absent, BACH1 redistributes to promoter-proximal regions and relocates toward canonical SATB1 sites. Critically, BACH1 facilitates the recruitment of cMAF, creating a BACH1-cMAF circuit that drives proinflammatory transcriptional programs in thymocytes. These pathogenic thymocyte states seed the periphery, where they imprint long-lasting gene expression changes in CD4⁺ T cells that resemble human SLE signatures. Genetic ablation of *Bach1* in *Satb1*-deficient T cells reverses these changes, normalizes cytokine levels, restores tissue integrity and prevents premature mortality.

Our findings uncover a previously unrecognized mechanism linking gene regulation to autoimmunity. By identifying BACH1 and cMAF as central effectors unleashed by SATB1 loss, we not only clarify the molecular basis of the autoimmune-like phenotype in *Satb1*-deficient mice but also nominate new therapeutic targets.

## Results

### BACH1 reprograms chromatin occupancy and inflammatory gene expression in *Satb1*-deficient thymocytes

Using DNA affinity chromatography coupled to mass spectrometry experiments^29,30^, to pull down proteins interacting with the *Rad50* DNAse I hypersensitive site 6 (RHS6) of the *Th2* locus control region (LCR)^31,32,33^ (151bp - the most conserved sequence between mouse and human genomes, participating in TH2-*Ifnγ* gene loci interchromosomal interactions) we have identified two proteins: SATB1 and BACH1^13^. We first performed immunofluorescence staining in murine wild-type (WT) thymocytes and observed that BACH1 displayed a subnuclear localization pattern similar to the previously described cage-like distribution of SATB1 protein (Fig. 1a). The two proteins showed extensive colocalization in the T cell nucleus (Fig. 1a).

**Fig. 1.**
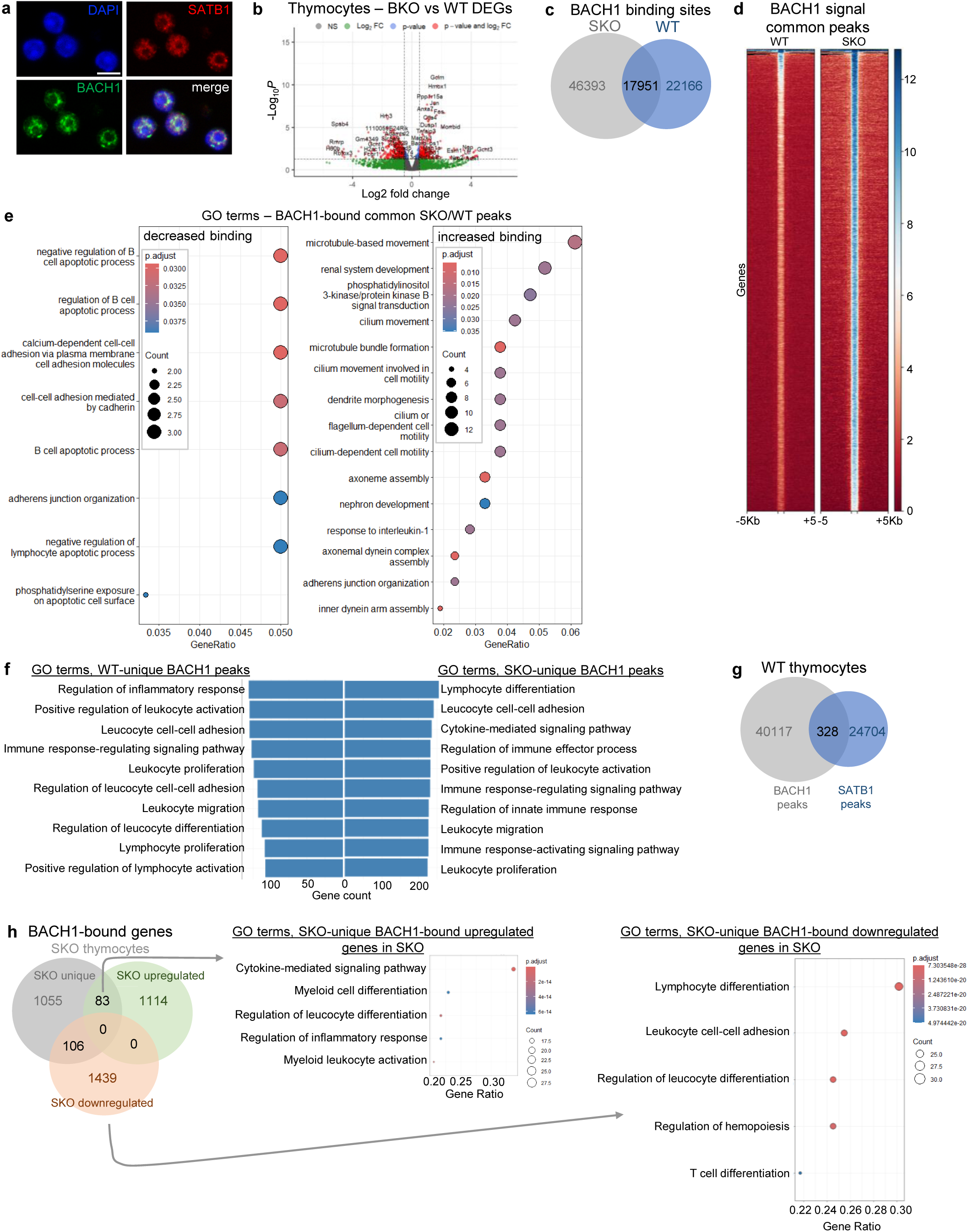
BACH1 reprograms chromatin occupancy and inflammatory gene expression in Satb1-deficient thymocytes. **a.** Immunofluorescence for BACH1 and SATB1 in primary C57BL/6 thymocytes. (Scale bar 5μm). **b.** Bulk RNAseq analysis and volcano plot of the differentially expressed genes (DEGs) in *Bach1*^−/-^ (BKO) versus WT thymocytes. 15.351 genes analyzed (Wald test, pvalue<0.05, fc>0.5). **c.** Venn diagram of BACH1 ChIPseq peaks in C57BL/6 (WT) and *Satb1^fl/fl^Cd4Cre* (SKO) thymocytes. **d.** Heatmap of BACH1 ChIPseq binding score at common BACH1 ChIP-seq peaks of WT and SKO thymocytes. (summit 1Kb ±5Kb). **E.** GO terms analysis of the BACH1-bound genes, in common BACH1 peaks between WT and SKO thymocytes (Hypergeometric test, FDR-adjusted pvalues, padj<0.05). **f.** GO terms of genes bound by BACH1 (ChIP-seq) uniquely in WT or SKO thymocytes (Hypergeometric test, FDR-adjusted pvalues, padj<0.05). **g.** Venn diagram of BACH1 ChIPseq and SATB1 HiChIP peaks in WT thymocytes. **h.** Venn diagram of the differentially expressed genes (bulk RNAseq) bound by BACH1 (ChIPseq) in SKO thymocytes. (top) GO Terms of the upregulated genes bound by BACH1 in SKO thymocytes. (bottom) GO Terms of the downregulated genes bound by BACH1 in SKO thymocytes (Hypergeometric test, FDR-adjusted pvalue, padj<0.05). *See also Extended Data Fig. 1 & 2*.

We and others have previously shown that conditional deletion of *Satb1* in T cells (driven by the *Cd4-Cre* transgene) results in a severe autoimmune-like phenotype in mice^10,11^. In the absence of SATB1, as deduced by the *Cd4Cre-Satb1*^fl/fl^ (SKO) thymocytes, the relative mRNA levels of *Bach1* were increased (Extended Data Fig. 1a). To examine whether BACH1 also plays an essential role in the development and differentiation of murine T cells, we used full body *Bach1*^−/-^ mice (hereafter referred to as BKO) and double knockout animals lacking both *Bach1* and *Satb1* genes in T cells (*Bach1*^−/-^/*Cd4Cre-Satb1*^fl/fl^, hereafter referred to as DKO). Antibodies directed against either the N-terminus (amino acids 133-513) or the C-terminus (amino acids 591-720) of BACH1 failed to detect BACH1 protein in *Bach1*^−/-^ (BKO) thymocytes (Extended Data Fig. 1b) or in *Cd4Cre*-*Satb1*^fl/fl^/*Bach1*^−/-^ (DKO) thymocytes (Extended Data Fig. 1c). In addition, co-immunoprecipitation experiments using murine thymocyte extracts did not detect any interaction between BACH1 and SATB1 proteins (Extended Data Fig. 1d).

To examine whether BACH1 also plays an essential role in the development and differentiation of murine T cells, we used *Bach1*^−/-^ (BKO) mice and we performed bulk RNA sequencing in thymocytes isolated from C57BL/6 and BKO mice. We identified 808 differentially expressed genes (294 upregulated and 514 downregulated) (Fig. 1b), supporting a role for BACH1 in the regulation of specific T-cell–mediated immune processes (Extended Data Fig. 1e). To map the BACH1 binding profile, we carried out chromatin immunoprecipitation followed by sequencing (ChIPseq) in wild type (WT) and SKO thymocytes. We identified 40.117 BACH1 binding sites in WT thymocytes and 64.344 sites in SKO thymocytes (Fig. 1c). Notably, BACH1 binding was enhanced in SKO compared with WT, at common genomic sites (Fig. 1d). Moreover, the newly emerged BACH1 peaks in SKO thymocytes were located closer to SATB1 binding sites (Extended Data Fig. 1f). BACH1 binding was also increased in promoter and promoter-proximal regions in SKO thymocytes (Extended Data Fig. 1g), with enhanced deposition at SATB1 binding sites (Extended Data Fig. 1h). These findings suggest that SATB1 acts as an access-regulator, delimiting BACH1 binding to discrete chromatin regions.

Gene ontology (GO) analysis of the WT/SKO BACH1-bound common peaks indicated increased binding in genes important for response to interleukin-1 in SKO thymocytes (Fig. 1e). WT-specific and SKO-specific BACH1 peaks were enriched in genes linked to leukocyte-specific pathways (Fig. 1f). Importantly, in SKO thymocytes, BACH1 binding near SATB1 sites was linked to genes involved in T-cell activation (Extended Data Fig. 1i,j). These findings suggest that SATB1 regulates the access to immune gene regions to safeguard proper spatiotemporal thymic development^34^. In SKO thymocytes, where SATB1 is absent, BACH1 binds regulatory elements near proinflammatory genes controlling leukocyte activation, immune response and T-cell selection, thereby driving inflammatory transcriptional programs.

Although immunofluorescence experiments indicated similar subnuclear localization for SATB1 and BACH1 in thymocytes, only a small fraction of overlapping peaks were detected in WT thymocytes by BACH1 ChIPseq and SATB1 HiChIP (328 common peaks; Fig. 1g).

The differentially expressed genes in SKO/WT bulk RNAseq that were bound by BACH1 in SKO thymocytes were enriched in pathways controlling T-cell differentiation and inflammatory immune responses (Fig. 1h). More specifically, the newly formed BACH1 peaks in SKO thymocytes showed increased deposition in the regulatory regions of 83 transcriptionally upregulated genes that are involved in cytokine-mediated signaling and effector processes (Fig. 1h). BACH1 can also bind enhancers of proinflammatory genes in both WT and SKO thymocytes; however, in WT cells, the repressive chromatin landscape enforced by SATB1 prevents their activation, whereas in SKO thymocytes, the absence of SATB1 allows the activation of these enhancers (Extended Data Fig. 2a,b). Taken together, these data highlight that upon SATB1 depletion, BACH1’s binding is unleashed and can mediate the expression of proinflammatory genes in developing thymocytes.

### Loss of BACH1 reverses the autoimmune-like SKO phenotype

Previous studies have shown that loss of SATB1 disrupts proper T-cell selection in the thymus, leading to the escape of autoreactive T cells into secondary lymphoid organs, followed by multi-organ infiltration and increased autoantibody production^35–37^. Our data now reveal crucial and opposing roles for SATB1 and BACH1 in developing thymocytes. Given the redistribution of BACH1 binding to immune gene loci in the absence of SATB1, we next asked whether BACH1 contributes to the autoimmune phenotype of SKO mice. To dissect these roles, we employed three knockout mouse models: *Cd4Cre-Satb1*^fl/fl^ (SKO), *Bach1*^−/-^ (BKO) and double *Cd4Cre*-*Satb1*^fl/fl^/*Bach1*^−/-^ (DKO).

Given that central tolerance defects often propagate to the periphery, we investigated whether SKO mice exhibited peripheral immune activation and whether this was reversed in the DKO mice. Strikingly, DKO mice completely lacked the extensive T cell infiltration observed in multiple tissues, including kidney, liver, pancreas, cornea and the lung of SKO mice, as revealed by hematoxylin and eosin (H&E) staining (Fig. 2a). This finding is consistent with prior reports that BKO mice display normal thymic T-cell development, intact peripheral lymphoid organs and a generally healthy phenotype^38^. By contrast, SKO and DKO mice both exhibited a developmental block at the CD4⁺CD8⁺ double-positive (DP) stage, as previously described for the loss of SATB1 in SKO mice^10^ (Fig. 2b and Extended Data Fig. 2c). Loss of BACH1 alone did not affect thymic T cell populations. Interestingly, DKO mice were more fertile (data not shown) and showed no overt phenotypic abnormalities compared with SKO mice.

**Fig. 2.**
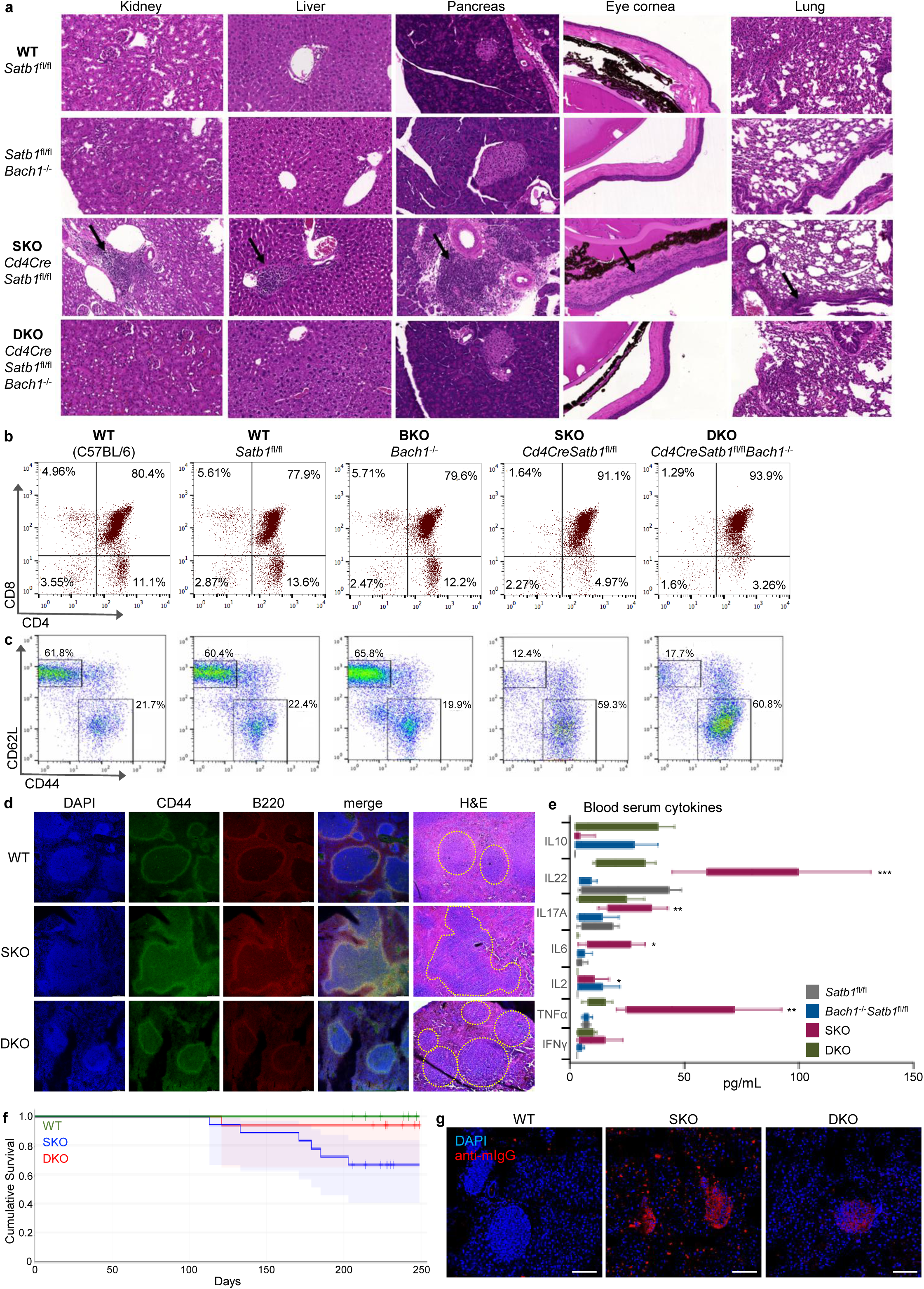
Loss of BACH1 reverses the autoimmune-like phenotype of *Satb1*-deficient mice. **a.** Hematoxylin and Eosin (H&E) staining of the indicated tissue sections from *Satb1*^fl/fl^ (WT), *Satb1^fl/fl^Bach1*^−/-^(BKO), *Cd4Cre*-*Satb1^fl/fl^* (SKO) and *Cd4Cre*-*Satb1^fl/fl^*/*Bach1*^−/-^ (DKO) mice. **b.** FACS analysis of thymocytes from C57BL/6 (WT), *Satb1^fl/fl^* (WT), *Bach1*^−/-^ (BKO), *Satb1^fl/fl^Cd4Cre* (SKO) and *Bach1*^−/-^/*Satb1^fl/fl^Cd4Cre* (DKO) for the expression of CD4 and CD8 markers. **c.** FACS analysis in secondary lymphoid organs (spleen and lymph nodes) of C57BL/6 (WT), *Satb1^fl/fl^* (WT), *Bach1*^−/-^ (BKO), *Satb1^fl/fl^Cd4Cre* (SKO), and *Bach1*^−/-^/*Satb1^fl/fl^Cd4Cre* (DKO) for the expression of CD44 and CD62L markers. **d.** Immunofluorescence experiments using anti-B220 (pan-B cell marker) and anti-CD44 (activated T cell marker) (7 repeats, WT n=4, SKO n=3, DKO n=5) as well as H&E staining in spleen sections of WT, SKO and DKO mice (4 repeats, WT n=3, SKO n=3, DKO n=4). **e.** Legendplex analysis for the cytokine profile in blood sera from WT (*Satb1^fl/fl^*), BKO, SKO and DKO mice (Welch’s t-test, n=8). **f.** Survival curves of male WT, BKO, SKO and DKO mice. Log-rank test p=0.014. (WT n=14, SKO n=18, DKO n=17). **g.** Autoantibody detection using WT, SKO and DKO sera on WT pancreas sections (3 repeats, 2 serum samples per genotype). *See also Extended Data Fig. 3*

To examine the activation state of peripheral T cells, we performed flow cytometry for CD44 (activation) and CD62L (naïve) markers in T cells from spleens and lymph nodes of WT, BKO, SKO and DKO mice. CD4⁺ T cells from SKO and DKO mice exhibited elevated CD44 marker expression, whereas BKO T cells resembled WT (Fig. 2c and Extended Data Fig. 2d). Because SKO mice displayed marked T cell activation, we next assessed splenic architecture. Immunofluorescence staining of B cells and activated T cells, combined with H&E staining, revealed disrupted splenic follicles and expansion of both T- and B-cell populations in SKO mice (Fig. 2d). This lymphoproliferative phenotype was accompanied by splenomegaly. By contrast, DKO mice exhibited splenic follicle morphology similar to WT, consistent with alleviation of the inflammatory phenotype.

To determine whether systemic inflammation was also ameliorated, we measured circulating cytokines using Legendplex multiplex analysis. SKO mice displayed elevated serum levels of proinflammatory cytokines, whereas these were markedly reduced in the DKO mice (Fig. 2e). This change correlated with increased survival of DKO animals compared with the premature mortality of SKO mice (Fig. 2f). Finally, consistent with the autoimmune phenotype, sera from SKO mice contained high levels of autoantibodies, while DKO sera lacked detectable autoantibodies (Fig. 2g).

Taken together, these data confirm that SKO mice develop a severe autoimmune-like phenotype, as previously reported^10^, whereas DKO mice - lacking both SATB1 and BACH1 in T cells - do not develop any notable autoimmune phenotype, although they show aberrant T cell development as in SKO. Thus, we assume that BACH1 plays an essential role in mediating the autoimmune features triggered by SATB1 loss.

### The pro inflammatory T, B and neutrophil cell populations of SKO spleens are lost in the DKO

SATB1 is a key factor safeguarding T cell development and activation, as its loss results in deregulated expression of proinflammatory genes, including members of chemokine gene clusters^39^. Because loss of BACH1 reverses most of the pathological features observed in the *Satb1*-deficient mice, we performed single-cell RNA sequencing (scRNAseq) of spleens from WT, SKO and DKO animals to delineate lineage-specific transcriptional programs that might drive the inflammatory phenotype in peripheral lymphoid tissues.

UMAP analysis revealed differences in both the number of cells (Fig. 3a) and composition (Fig. 3b, Extended Data Fig. 3a) of splenic cell clusters across genotypes. SKO spleens showed a great loss of cells in cluster 0, which was partially restored in DKO and an expansion of clusters 1–6, which were largely reduced in DKO. Cluster 0 consisted mainly of epithelial cells, stromal cells, endothelial cells and fibroblasts and its absence in SKO, correlated with structural defects in splenic follicles. By contrast, loss of SATB1 led to an expansion of T, B and neutrophil populations, consistent with immune activation and inflammation. Strikingly, these features were reversed in DKO spleens.

**Fig. 3.**
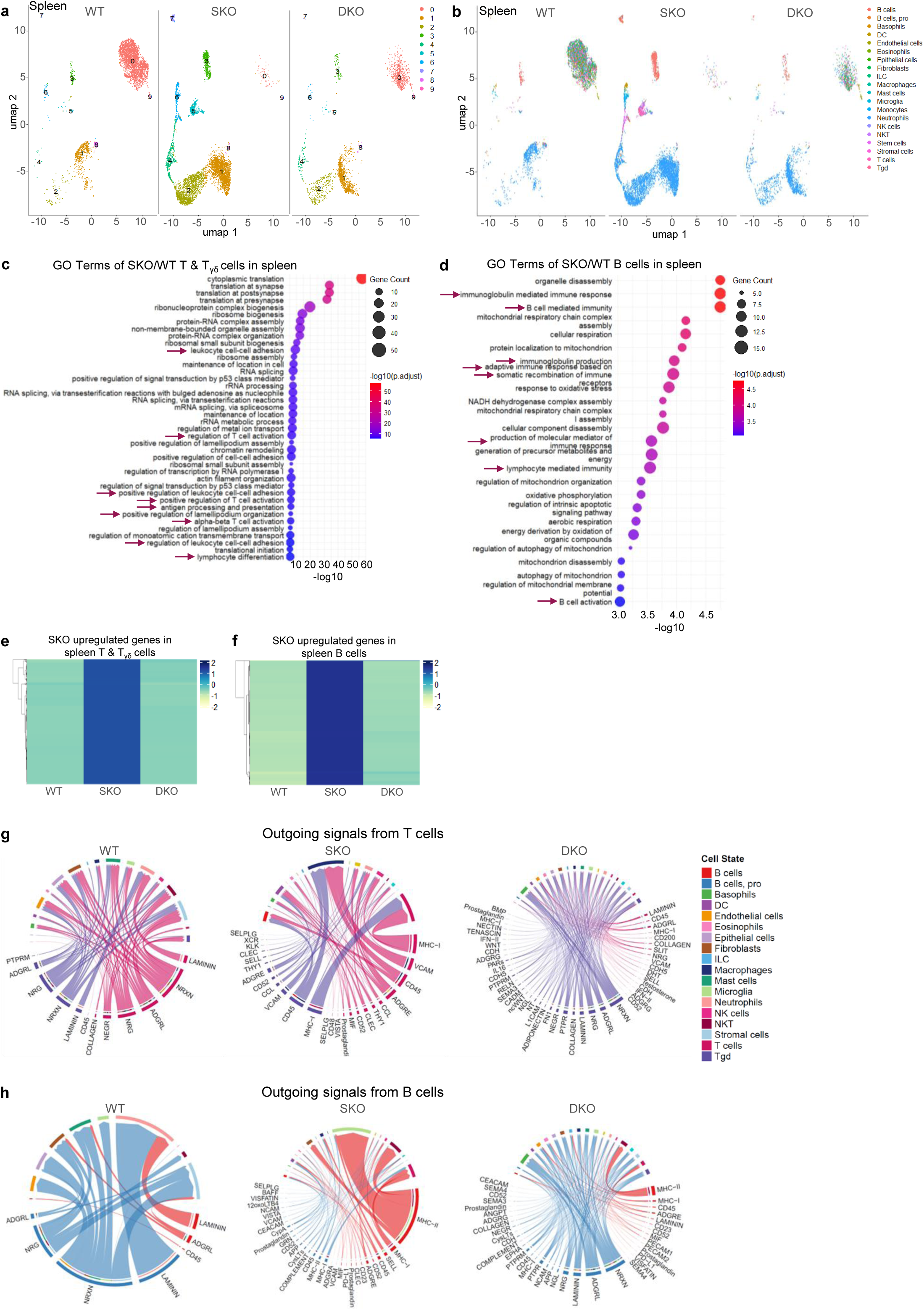
The pro inflammatory T, B and neutrophil cell populations of SKO spleens are lost in the DKO. **a, b.** UMAP of splenic scRNA-seq data split either by cluster(A) or cell type (B) from wild type, SKO and DKO mice. **c.** GO Terms of the upregulated genes of SKO/WT differential gene expression analysis in T cells and Tγδ cells, from spleen scRNAseq data (Hypergeometric test, FDR-adjusted pvalue, padj<0.05). **d.** GO Terms of the upregulated genes of SKO/WT differential gene expression analysis in B cells from spleen scRNAseq data (Hypergeometric test, FDR-adjusted pvalue, padj<0.05). **e.** Heatmap of the relative expression levels of SKO upregulated genes in WT, SKO and DKO T cells and Tγδ cells from spleen scRNAseq (Wald test, pvalue<0.05, fc>0.5). **f.** Heatmap of the relative expression levels of SKO upregulated genes in WT, SKO and DKO B cells from spleen scRNAseq (Wald test, pvalue<0.05, fc>0.5). **g.** Cellchat analysis of ligand-receptor interaction probability of all outgoing signals from T cells in WT, SKO and DKO mice from spleen scRNAseq data. **h.** Cellchat analysis of ligand-receptor interaction probability of all outgoing signals from B cells in WT, SKO and DKO mice from spleen scRNAseq data. *See also Extended Data Fig. 3*.

Differential gene expression analysis of T cell populations showed increased expression of genes involved in activation, adhesion and proliferation (Fig. 3c and Extended Data Fig. 3b). In turn, B cells displayed elevated expression of genes linked to MHC complex assembly, antigen processing and presentation and B-cell–mediated immunity (Fig. 3d and Extended Data Fig. 3c). These findings point towards the possibility that ectopically activated T cells in SKO spleens aberrantly stimulate B cells, driving immunoglobulin production and activation of humoral responses. This is consistent with our observation of increased autoantibodies in SKO sera. Importantly, the expression levels of the proinflammatory genes in T and B cells of SKO mice - including pathways for immunoglobulin-mediated immunity, B-cell activation and immune responses - were ameliorated in DKO mice (Fig. 3e, f).

To further dissect the intercellular communication, we performed CellChat analysis on the spleen scRNAseq data. We found that SKO T cells were predicted to have stronger ligand-receptor interactions with macrophages, B cells, dendritic cells and neutrophils, particularly via MHC-I, MHC-II, CD45 and CCL signaling (Fig. 3g). These signals were absent or greatly reduced in spleen DKO T cells. Outgoing signals from the SKO T cells to B cells and neutrophils included increased CD45 signaling, as well as heightened CCL (CC chemokine ligands, when bound to chemokine receptors drive leukocyte chemotaxis, activation and tissue recruitment) and ADGRE (Adhesion G-protein-coupled receptor E family, promote cell-cell adhesion, migration and immunologic synapse stability) signaling to neutrophils - all diminished in the DKO T cells (Extended Data Fig. 3d, upper panel). In parallel, B cells from SKO spleens exhibited increased prediction of ligand-receptor communication with T cells, NKT cells and basophils through CD45, MHC-I and MHC-II molecules (Fig. 3h and Extended Data Fig. 3d, lower panel), whereas these predicted interactions were lost or strongly reduced in DKO splenic B cells.

Together, these results demonstrate that loss of SATB1 drives the expansion and activation of pro-inflammatory T, B and neutrophil populations and enhances their crosstalk within the spleen to support their activation. Importantly, loss of BACH1 in the SKO background reverses these proinflammatory signatures and restores cellular communication toward a WT-like state.

### BACH1 cooperates with cMAF in the newly formed proinflammatory cell populations of SKO thymi

The autoimmune-like phenotype manifested in peripheral lymphoid organs and multiple tissues is caused by the thymus-specific loss of *Satb1* early during T cell development, at the CD4⁺CD8⁺ double-positive (DP) stage^10,39,40^. To identify the pathogenic thymocyte populations responsible for driving this phenotype in SKO mice, we performed scRNAseq on whole thymi from WT, SKO and DKO animals. UMAP analysis revealed that loss of SATB1 resulted in pronounced changes of several clusters (Fig. 4a, and Extended Data Fig. 4). Clusters 2, 4 and 7 were massively expanded in SKO thymi, whereas clusters 5 and 6 were reduced. Differential gene expression and enrichment analyses of clusters 1 and 3 in SKO versus WT indicated increased expression of genes related to cytokine receptor activity, type II interferon production and immune receptor signaling (Extended Data Fig. 5a, b). Although these clusters persisted in DKO thymi, their gene expression profiles were downregulated to WT levels. Among the upregulated proinflammatory genes were *Ifngr1*, *Gzma*, *Il18r1*, *Il7r* and *Gata3*, whereas critical developmental genes such as *Tcf7*, *Ly6d*, *Themis* and *Arpp21* were downregulated. Clusters 2, 4 and 7, which were expanded in SKO, were absent in DKO thymi. Gene set enrichment analysis (GSEA) indicated that in SKO thymi these clusters showed enrichment for oxidative phosphorylation, fatty acid metabolism and glycolysis, therefore metabolically active cells consistent with their functional activation, all of which were reversed in DKO (Extended Data Fig. 5c-e). Mitotracker staining confirmed increased mitochondrial load in SKO thymocytes, which was normalized in DKO (Fig. 4b). These data indicate that SATB1 loss drives a metabolic shift in thymocytes toward increased oxidative and glycolytic flux consistent with activation and proliferation programs. *Bach1* deletion, in DKO thymi, restores a quiescent metabolic state, implicating BACH1 in sustaining this metabolic rewiring.

**Fig. 4.**
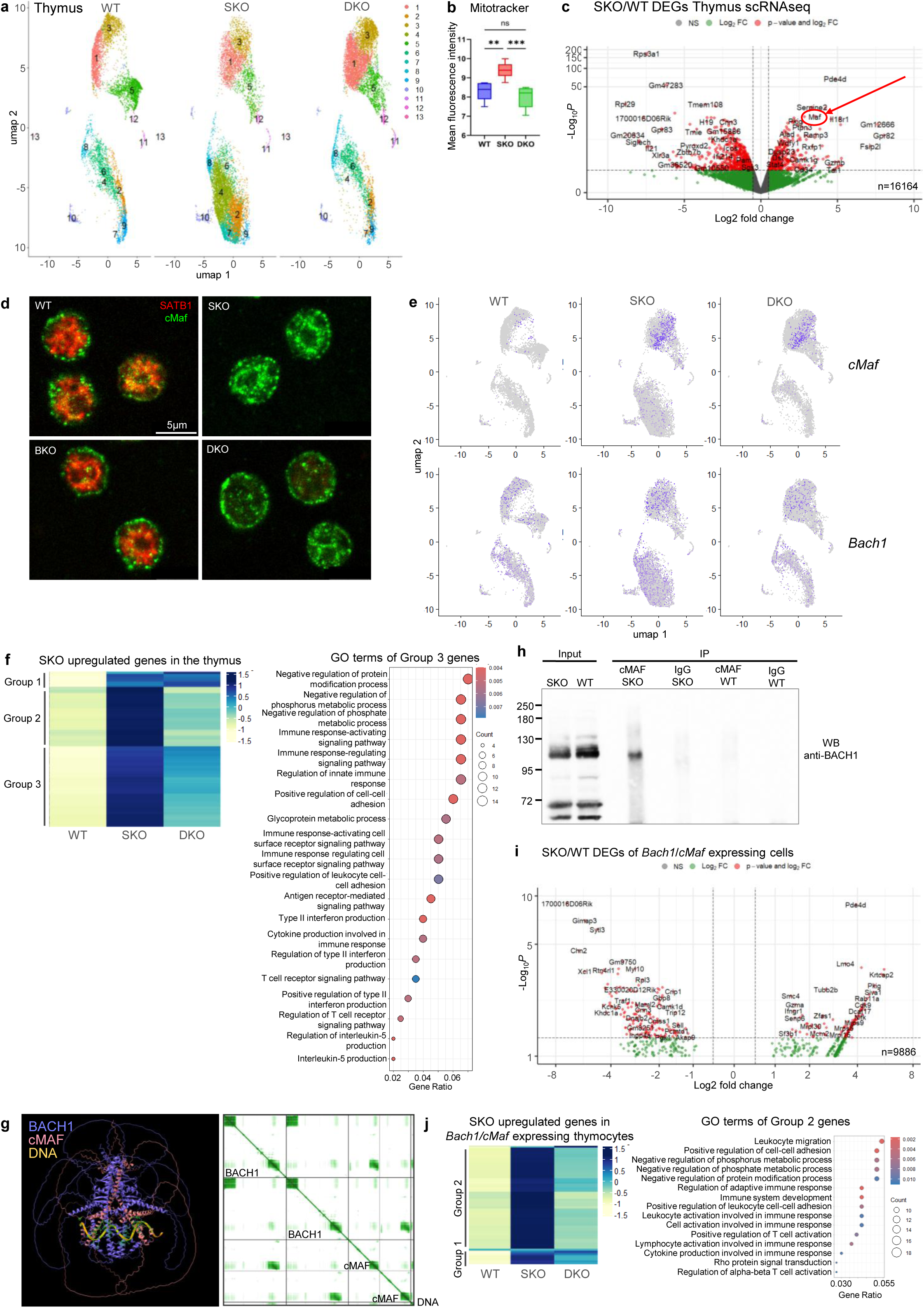
Loss of SATB1 enables BACH1/cMAF-driven proinflammatory programs in thymocytes. **a.** UMAP visualization of whole thymi scRNAseq split by cluster from WT, SKO and DKO mice. **b.** Quantification of mean fluorescence intensity of Mitotracker immunofluorescence in WT, SKO and DKO thymocytes (n=3). **c.** Volcano plot of the differential gene expression analysis after pseudobulking of whole thymi scRNAseq data in SKO/WT thymocytes (Wald test, pvalue<0.05, fc>0.5). **d.** Immunofluorescence analysis for SATB1 and cMAF expression in thymocytes from WT, BKO, SKO and DKO mice (n=5). **e.** UMAP visualization of whole thymi scRNAseq data highlighting *cMaf* and *Bach1* expression in WT, SKO and DKO thymocytes. **f.** Heatmap of the relative gene expression levels of SKO upregulated genes from (C) whole thymi scRNAseq data in WT, SKO and DKO thymocytes. **g.** AlphaFold modeling for the prediction of BACH1 interaction with cMAF and DNA. **h**. Co-immunoprecipitation of BACH1 and cMAF from total murine thymocyte protein extracts (10% input loaded, n=4). **i.** Volcano plot for the differentially expressed genes of *Bach1*/*cMaf*-expressing cells from thymi scRNAseq data in SKO/WT (Wald test, pvalue<0.05, fc>0.5. **j.** Heatmap of the relative gene expression levels for the SKO upregulated genes in *Bach1*/*cMaf* expressing cells from thymi scRNAseq data in WT, SKO and DKO thymocytes from (H). *See also* Extended Data Fig. 4-6

Differential gene expression analysis further revealed that *cMaf*, encoding the transcription factor cMAF, was one of the most upregulated genes in SKO thymocytes (Fig. 4c). cMAF is a versatile regulator of T-cell biology, promoting Th17 and Tfh differentiation while also contributing to regulatory programs, depending on context^21,22,28^. Bulk RNA-seq confirmed that *cMaf* was highly upregulated in SKO and DKO compared to WT thymocytes (Extended Data Fig. 6a). H3K27ac HiChIP data showed increased promoter-enhancer loops at the *cMaf* locus in SKO thymocytes (Extended Data Fig. 6b), consistent with enhanced expression, indicating that *cMaf* expression is dependent on chromatin reorganization of the locus. Increased *cMaf* mRNA levels were validated by RT-qPCR (Extended Data Fig. 6c) and elevated protein levels by western blot analysis (Extended Data Fig. 6d).

Immunofluorescence experiments revealed that cMAF protein mainly localized to the nucleus with a cage-like distribution in SKO thymocytes, similar to SATB1 and BACH1 proteins (Fig. 4d). Nuclear localization of cMAF was highest in SKO thymocytes and reduced in DKO (Extended Data Fig. 6e). scRNAseq showed that *cMaf* expression was concentrated in clusters 1 and 3 (Fig. 4e). Although these clusters persisted in DKO, *cMaf* and other proinflammatory transcripts were expressed at lower levels compared to SKO (Fig. 4f). Cell identity analysis indicated that *cMaf*-expressing cells corresponded largely to immature single-positive (ISP) T cells, which were expanded in SKO but normalized in DKO thymocytes (Extended Data Fig. 6f). These findings suggest a developmental blockade at the ISP stage, accompanied by increased *cMaf* expression, in SKO thymocytes that is alleviated in DKO mice.

To test whether cMAF interacts with BACH1, we first performed AlphaFold structural predictions, which indicated hetero-oligomerization via their bZIP domains, forming a tetramer (two BACH1 and two cMAF molecules) stabilized on DNA (Fig. 4g and Extended Data Fig. 6g). This arrangement is consistent with structural studies of BACH1-small MAF interactions^41^. Co-immunoprecipitation of thymocyte extracts confirmed the interaction between BACH1 and cMAF in SKO thymocytes (Fig. 4h and Extended Data Fig. 6h). Immunofluorescence analysis of spleen sections further revealed that while BACH1 and cMAF were expressed in distinct follicular populations in WT, they were co-expressed in disrupted follicles of SKO spleens (Extended Data Fig. 6i).

Finally, differential gene expression analysis of *Bach1*/*cMaf* co-expressing thymocytes from thymi scRNAseq data showed increased expression of genes involved in T cell-mediated immunity, adaptive immune responses and cytokine receptor signaling in SKO, compared to WT mice (Fig. 4i). Many of these proinflammatory genes were downregulated in DKO thymocytes (Fig. 4j).

Together, these data indicate that in the absence of SATB1, cMAF is upregulated, localizes to the nucleus, and cooperates with BACH1 to drive pathogenic transcriptional programs in thymocytes. Loss of BACH1 prevents cMAF from maintaining this proinflammatory state, thereby alleviating the autoimmune-like phenotype.

### cMAF plays an important role in the gene regulatory network of the autoimmune-like phenotype

Although cMAF has been previously shown to excerpt an important anti-inflammatory effect^42^, its role in promoting the progression of autoimmune disorders^26^ or the activation of inflammatory responses^27^ is also profound. To assess the role of cMAF in the gene regulatory networks of developing T cells, we performed ChIPseq experiments against cMAF in SKO and DKO thymocytes, where it is transcriptionaly upregulated in the absence of SATB1. ChIPseq analysis revealed a dramatic decrease of cMAF binding in DKO thymocytes, with 59.902 peaks in SKO reduced to 6.576 in DKO thymocytes (Fig. 5a). This indicates that BACH1 is required for the widespread genomic binding of cMAF. Even at the subset of sites bound by cMAF in both the SKO and DKO thymocytes, binding was markedly weaker in DKO thymocytes. We analyzed the signal strength of cMAF binding in the common binding sites we have detected in the SKO and DKO thymocytes and we found that in the SKO thymocytes, where SATB1 is absent, cMAF binds stronger compared to the weaker signal strength detected in DKO thymocytes, where both SATB1 and BACH1 are absent (Fig. 5b). The total cMAF peaks annotation indicated that the binding sites of cMAF in the SKO thymocytes were enriched for genes involved in lymphocyte differentiation, proliferation, cytokine-mediated and immune response-activating signaling pathways, while in DKO thymocytes the gene enrichment analysis indicated genes involved in more general cellular and developmental processes (Fig. 5c). SKO-unique cMAF peaks were enriched for genes linked to lymphocyte activation and differentiation (Extended Data Fig. 7a). These genes formed immune-related networks and included many that were upregulated in SKO thymocytes (Extended Data Fig. 7b, c), indicating that cMAF binding in the absence of SATB1 is directed toward proinflammatory programs. These findings indicate that cMAF binds chromatin close to proinflammatory genes only in SKO thymocytes, in the absence of SATB1 and the concomitant presence of BACH1.

**Fig. 5.**
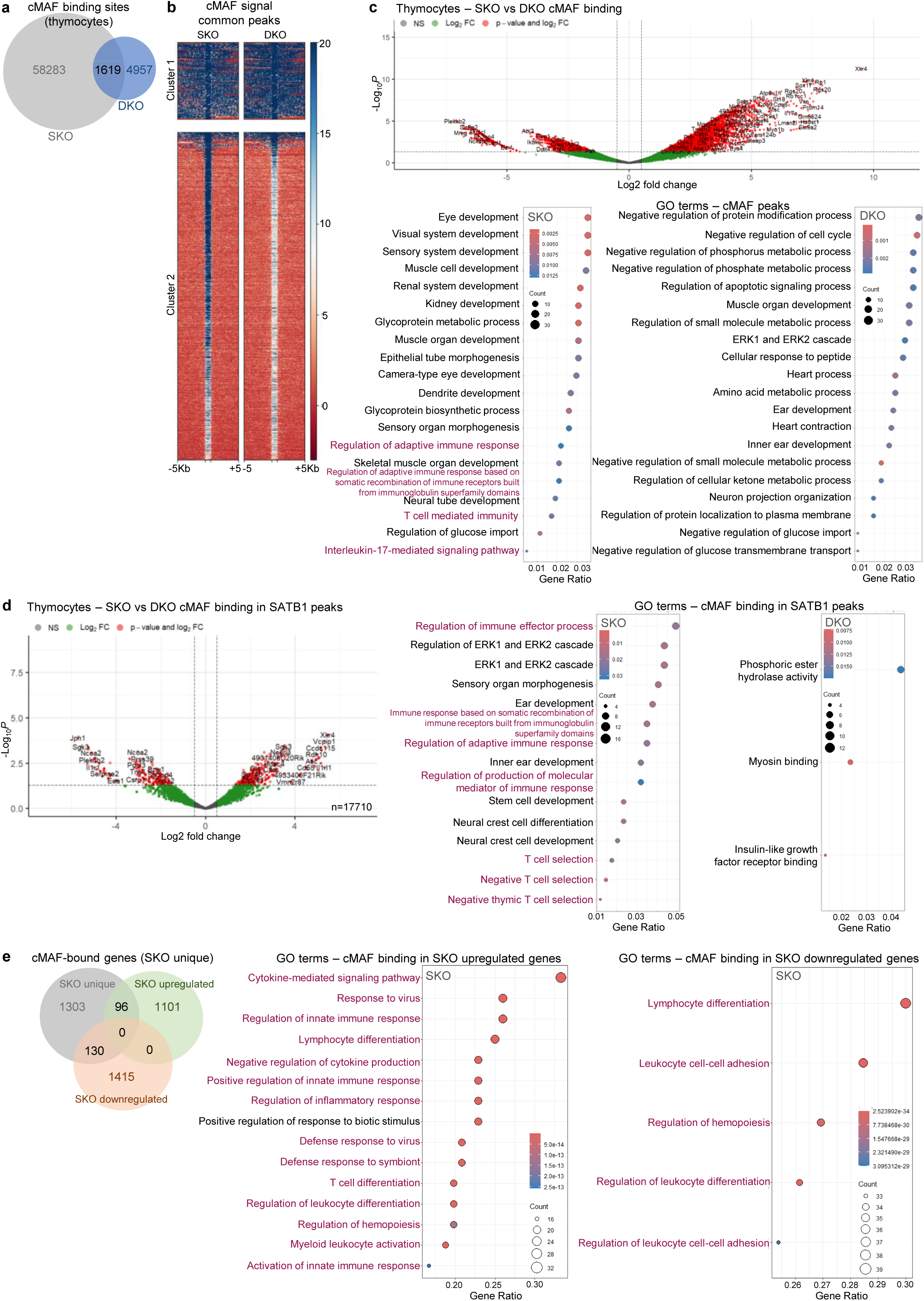
BACH1 enables cMAF binding to proinflammatory loci in thymocytes. **a.** Venn diagram for the overlap of cMAF ChIPseq binding sites in SKO and DKO thymocytes. **b.** K-mean clustering and heatmap of cMAF ChIP-seq binding score at common cMAF ChIPseq peaks of SKO and DKO thymocytes (summit 1Kb ±5Kb). **c.** Volcano plot of the differential binding of cMAF at ChIPseq peaks of SKO and DKO thymocytes (Exact test, pvalue<0.05, fc>0.5). **d.** Volcano plot of the differential binding analysis of cMAF at SATB1 peaks in SKO and DKO thymocytes (Exact test, pvalue<0.05, fc>0.5) and enriched GO terms of the bound genes (Hypergeometric test, FDR-adjusted pvalue, padj<0.05). **e.** Venn diagram and enriched GO terms of the differentially expressed genes (bulk RNAseq) bound by cMAF in SKO thymocytes (Hypergeometric test, FDR-adjusted pvalue, padj<0.05). *See also Extended Data Fig. 7*.

Since *cMaf* is greatly upregulated in the absence of SATB1, we then wanted to check whether in SKO and DKO thymocytes cMAF can bind to regulatory regions of genes regulated by SATB1. Therefore, we have checked for cMAF binding in SATB1 peaks in SKO and DKO thymocytes and found that in SKO thymocytes cMAF, similar to BACH1, was bound to genes involved in the regulation of the adaptive immune response (Fig. 5d). More specifically, cMAF was bound to the regulatory regions of 96 genes that were upregulated in SKO versus WT thymocytes from bulkRNA-seq data (Fig. 5e). More importantly, these differentially expressed genes in SKO thymocytes, bound by cMAF, are involved in the regulation of inflammatory immune responses and the aberrant activation of T cells in SKO thymocytes. In contrary, the genes bound by cMAF uniquely in DKO thymocytes or the common cMAF-bound genes in SKO and DKO thymocytes regulate processes other than the activation and differentiation of the immune system (Extended Data Fig. 7d, e).

These finding indicate that *cMaf* expression is de-repressed in the absence of SATB1 and then its protein product is recruited to genomic regions that regulate the development and activation of thymocytes. Although cMAF is expressed in DKO thymocytes, it displays reduced nuclear localization and fails to bind to genomic targets, consistent with a requirement for BACH1 in stabilizing its chromatin association.

### BACH1 and cMAF coregulate a proinflammatory gene network upregulated in the absence of SATB1

To delineate the impact of cooperative BACH1 and cMAF binding on proinflammatory genes in the absence of SATB1, we integrated the data from ChIPseq experiments for both factors. Since BACH1 is expressed in WT thymocytes and loss of BACH1 in DKO thymocytes results in a massive reduction of cMAF chromatin binding (Fig. 5b), we performed k-means clustering to assess cMAF occupancy at BACH1 peaks that were either shared between WT and SKO thymocytes (Fig. 6a) or uniquely found in SKO thymocytes (Fig. 6b). In common BACH1 peaks, cMAF signal score was significantly stronger in SKO than in DKO thymocytes, particularly in cluster 1, which was enriched for genes involved in cytokine-mediated leukocyte regulation (Fig. 6a, middle panel). By contrast, cluster 2 comprised regions with stronger cMAF binding in DKO and was enriched for genes associated with interleukin-10 signaling, consistent with the anti-inflammatory role of this cytokine and the reversal of autoimmune-like phenotype observed in DKO compared with SKO mice (Fig. 6a, far right panel). When we analyzed cMAF binding at SKO-unique BACH1 peaks, we identified four clusters with distinct patterns: clusters 1 and 2 displayed strong cMAF binding in SKO, cluster 3 showed stronger binding in DKO and cluster 4 displayed weak cMAF occupancy overall (Fig. 6b). Functional annotation revealed that genes of clusters 1 and 2 were enriched in T cell activation pathways (Fig. 6b, middle panel), whereas cluster 3 genes were linked to the negative regulation of inflammatory responses (Fig. 6b, far right panel). These observations align with the ameliorated inflammatory phenotype of DKO mice.

**Fig. 6.**
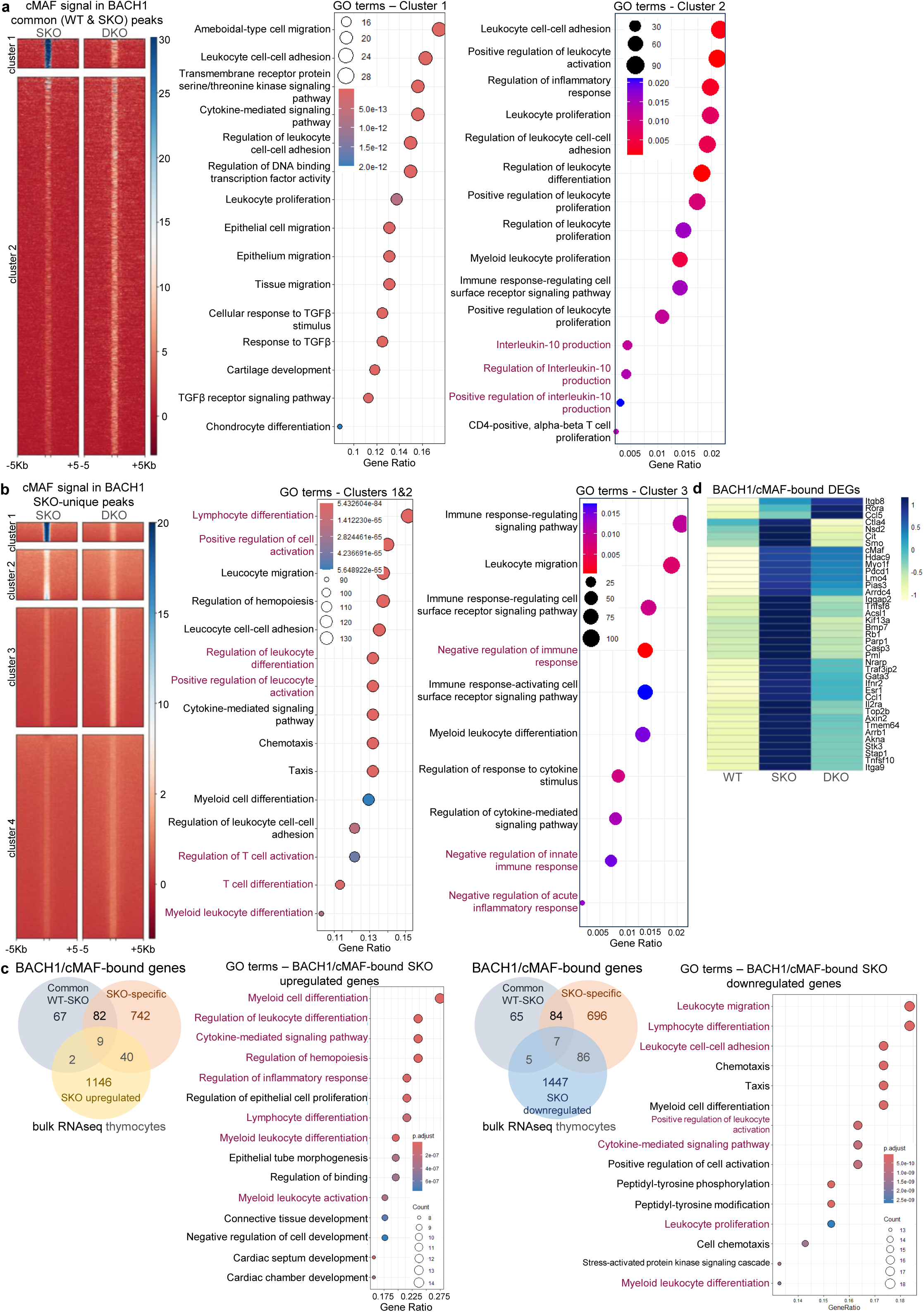
Cooperative binding of BACH1 and cMAF drives proinflammatory gene expression in *Satb1*-deficient thymocytes. **a.** K-means clustering and heatmap of cMAF ChIPseq signal score in WT/SKO common BACH1 peaks (summit 1Kb ±5Kb) and gene ontology of the bound genomic coordinates in SKO and DKO thymocytes of clusters 1 and 2 (Hypergeometric test, FDR-adjusted pvalue, padj<0.05). **b.** K-means clustering and heatmap of cMAF ChIPseq signal score in SKO newly formed BACH1 peaks (summit 1Kb ±5Kb) and gene ontology of the bound genomic coordinates in SKO and DKO thymocytes of clusters 1, 2 and 3 (Hypergeometric test, FDR-adjusted pvalue, padj<0.05). **c.** Venn diagram and gene ontology of the differentially expressed genes (bulk RNA-seq) bound by cMAF in the WT/SKO common or SKO newly formed BACH1 peaks (Hypergeometric test, FDR-adjusted pvalue, padj<0.05). **d.** Heatmap of the relative gene expression levels for the SKO upregulated genes (thymus scRNAseq) in WT, SKO and DKO thymocytes that are bound by BACH1/cMAF in SKO thymocytes (Hypergeometric test, FDR-adjusted pvalue, padj<0.05). *See also Extended Data Fig. 8*.

We next tested whether concomitant BACH1 and cMAF binding at common or SKO-specific BACH1 peaks impacted transcriptional programs in thymocytes. Bulk RNAseq analysis identified 51 genes that were bound by both BACH1 and cMAF and were upregulated in SKO compared to WT thymocytes (Fig. 6c). These genes are involved in leukocyte differentiation, activation and inflammatory responses. Similar results emerged by the intersection of the BACH1/cMAF co-bound genes with the upregulated genes in the scRNAseq analysis of SKO thymocytes, which are are in processes such as T-cell activation and differentiation pathways (Extended Data Fig. 8a). Notably, downregulated genes bound by BACH1 and cMAF in SKO thymocytes were also involved in leukocyte differentiation and activation (Fig. 6c, far right panel). Importantly, the increased expression levels of most of these 51 genes (36/51) in SKO thymocytes were decreased in DKO thymocytes (Fig. 6d). These findings highlight that BACH1-cMAF cooperation is essential for activating a proinflammatory gene network in the absence of SATB1, driving the pathogenic potential of SKO thymocytes.

### BACH1 cooperation with cMAF regulates the gene expression network of peripheral CD4^+^ T cells in SKO mice, leading to human autoimmune-like pathophysiology

Beyond the proinflammatory genes co-bound by BACH1 and cMAF during thymocyte development, we identified additional target genes whose expression was unaffected by SATB1 loss in the thymus but were strongly occupied by BACH1 and cMAF. To determine whether these genes were transcriptionally activated after T cell maturation, we performed bulk RNAseq of peripheral (splenic) CD4⁺ T cells from WT and SKO mice. Differential expression analysis revealed the transcriptional upregulation of genes associated with cytokine-mediated inflammatory responses in SKO compared with WT CD4⁺ T cells (Fig. 7a).

**Fig. 7.**
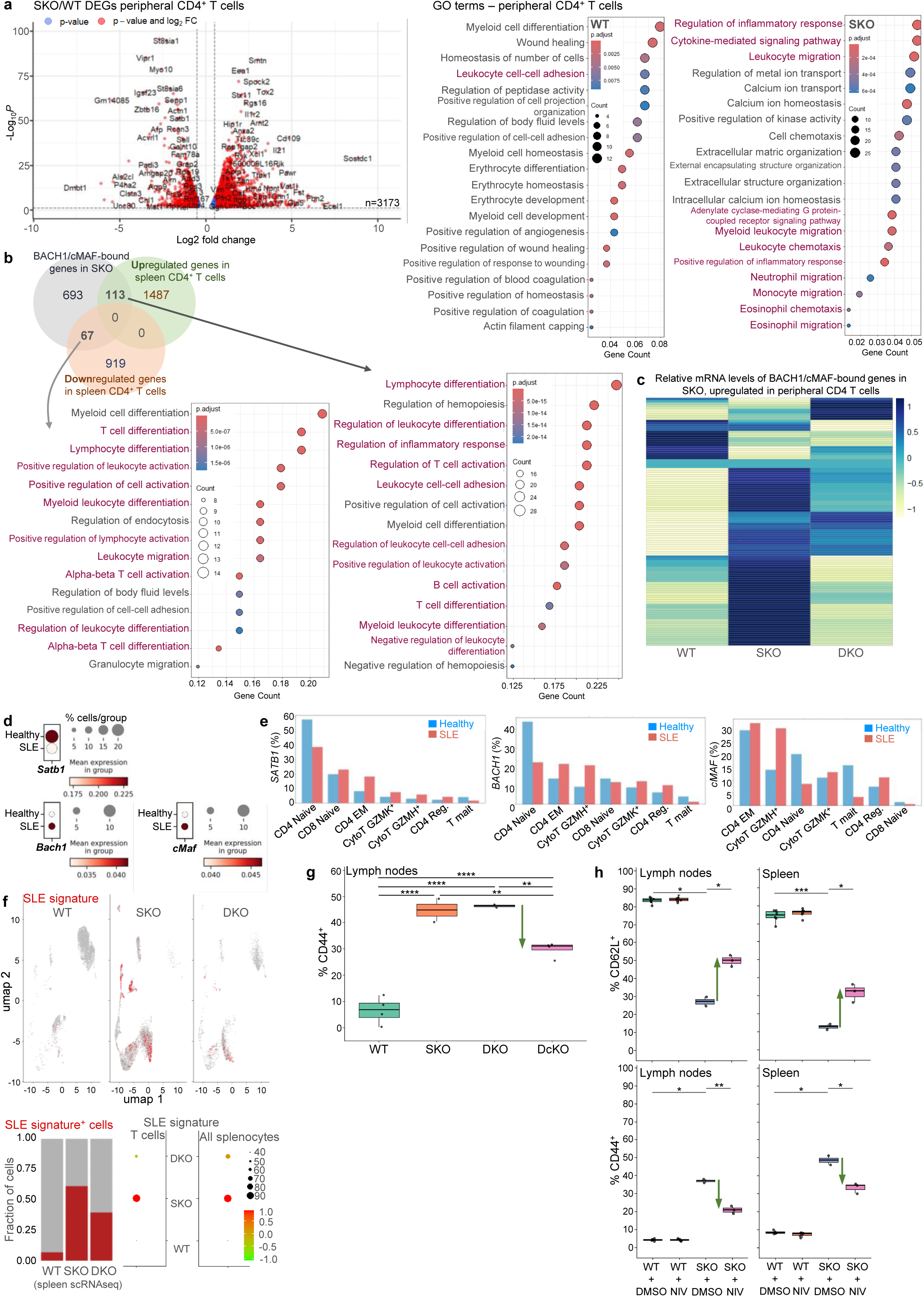
Peripheral CD4⁺ T cells in *Satb1*-deficient mice acquire BACH1/cMAF-driven SLE-like signatures. **a.** Volcano Plot and gene ontology of the differential gene expression analysis of bulk RNAseq data in SKO/WT peripheral CD4^+^ T cells. (Wald test, pvalue<0.05, fc>0.5) (Hypergeometric test, FDR-adjusted pvalue, padj<0.05). **b.** Venn diagram and gene ontology of the differentially expressed genes in peripheral SKO/WT CD4^+^ T cells (bulk RNAseq) that are bound by BACH1/cMAF in SKO thymocytes (Hypergeometric test, FDR-adjusted pvalue, padj<0.05). **c.** Heatmap of the relative gene expression levels of the SKO upregulated genes (spleen scRNAseq) in WT, SKO and DKO peripheral T cells that are bound by BACH1/cMAF in SKO thymocytes. **d.** Dotplot of the relative expression levels of *SATB1*, *cMAF* and *BACH1* in T cells of healthy individuals and SLE patients from scRNAseq data of PBMCs. **e.** Barplot of the relative gene expression levels of *SATB1*, *BACH1* and *cMAF*, in T cell populations of healthy individuals and SLE patients from scRNAseq data of PBMCs. **f.** UMAP visualization of the enrichment score of the SLE T cell signature in WT, SKO and DKO spleen scRNAseq data (Pairwise comparisons, Wilcoxon rank sum test WTvsSKO: pvalue<2e-16, SKOvsDKO: pvalue=0.0037). A red dot indicates a cell with SLE signature. **g.** FACS analysis of CD44^+^ cells from lymph nodes of WT, SKO, DKO and double conditional knockout cells for *Satb1* and *cMaf* (DcKO) mice. **h.** FACS analysis of CD62L^+^ and CD44^+^ cells from peripheral lymphoid organs (lymph nodes, spleen) in WT (DMSO or Nivalenol-treated) and SKO (DMSO and Nivalenol-treated) mice. *See also Extended Data Fig. 7*.

We next wanted to test whether these differentially expressed genes that were involved in promoting inflammatory immune responses in SKO CD4^+^ T cells were those demonstrating co-binding of BACH1 and cMAF. For this, we intersected the BACH1/cMAF-bound genes in thymocytes with the differentially expressed genes in splenic CD4⁺ T cells. This analysis identified 113 genes that were bound by BACH1 and cMAF in thymocytes and were upregulated in SKO splenic CD4^+^ T cells, many of which are involved in T-cell activation, differentiation and inflammatory responses. An additional 67 genes bound by BACH1 and cMAF were downregulated in SKO splenic CD4^+^ T cells and were likewise enriched in leukocyte activation and differentiation pathways (Fig. 7b). Notably, these 113 genes were bound early during thymocyte development but became upregulated only after T cell exit into the periphery. scRNAseq in WT, SKO and DKO spleens confirmed that the increased expression levels of most of these genes in SKO CD4⁺ T cells were reversed in DKO CD4⁺ T cells (Fig. 7c).

To further assess whether these networks were associated with enhancer activity, we performed H3K4me1^43–47^ ChIP-seq in WT, SKO and DKO thymocytes. We observed strong BACH1 and cMAF binding at genomic regions marked by H3K4me1 in all genotypes, with particularly high levels in SKO thymocytes (Extended Data Fig. 8b). BACH1/cMAF bound H3K4me1 regions in SKO thymocytes corresponded to 81 upregulated and 46 downregulated genes (bulk RNAseq data) of SKO splenic CD4^+^ T cells. (Extended Data Fig. 8c). Both gene sets were associated with T cell activation and differentiation. Importantly, 93% of the upregulated genes marked by H3K4me1 and bound by BACH1/cMAF in SKO showed decreased expression in DKO splenic T cells (scRNAseq data), underscoring the role of BACH1/cMAF in priming enhancer regions of proinflammatory genes (Extended Data Fig. 8d).

Quite importantly, to test the relevance of these findings to human autoimmunity, we analyzed scRNAseq datasets from peripheral blood mononuclear cells (PBMCs) of healthy individuals and systemic lupus erythematosus (SLE) patients^48^ (Extended Data Fig. 8e). SATB1 expression was downregulated, whereas BACH1 and cMAF were upregulated in SLE T cells, paralleling our mouse SKO model (Fig. 7d). Analysis of T cell subsets showed SATB1 enrichment in naïve CD4⁺ and CD8⁺ T cells in both cohorts, while BACH1 and cMAF were preferentially expressed in effector memory CD4⁺ T cells and cytotoxic GZMH⁺ T cells in SLE patients - populations associated with pathogenic inflammation (Fig. 7e). *BACH1* was expressed in 23% of naïve CD4 T cells and was also expressed in high levels in effector memory CD4 T cells (22%) and cytotoxic GZMH^+^ T cells (21%), populations that are characterized by a proinflammatory gene signature and are expanded during the progression of the disease. Additionally, *cMAF* was mainly expressed in effector memory CD4^+^ T cells (28%) and cytotoxic GZMH^+^ T cells (31%), indicating a similar increased expression in proinflammatory T cell populations in SLE patients.

We next examined whether the SLE inflammatory signature (100 genes overexpressed in SLE T cells) overlapped with the gene networks identified in SKO mice. Gene set scoring analysis in spleen scRNAseq experiments demonstrated that the SLE signature was strongly enriched in the splenic SKO T cells (Fig. 7f, upper panel). The SLE signature was detected in 61.2% of SKO splenocytes, including T cells, neutrophils and basophils, but dropped to 39.6% in DKO splenocytes (Fig. 7f, lower panel).

Finally, to test whether cMAF itself is necessary for driving the SKO autoimmune phenotype, we crossed *Cd4*Cre*Satb1*^fl/lf^ mice with *cMaf*^fl/fl^ animals to generate *Cd4*Cre*Satb1*^fl/lf^*Maf*^fl/fl^ double conditional knockout (DcKO) mice. DcKO animals exhibited normal development, fertility and survival compared to SKO littermates (data not shown). Importantly, lymph node CD4^+^ T cells from DcKO mice displayed reduced frequencies of CD44⁺ effector memory T cells compared to SKO mice (Fig. 7g). Similarly, the pharmacological inhibition of cMAF, using the small-molecule inhibitor Nivalenol, led to the reduced frequency of effector memory CD44⁺ T cells, in spleens and lymph nodes (Fig. 7h, lower panels), the concomitant increase in the frequency of naïve CD4+ T cells (Fig. 7h, upper panels) and the decreased expression of proinflammatory cMAF target genes (*Ifng, Ccl5, Il12rb2, Cd5l*) in SKO thymocytes (Extended Data Fig. 8f).

Together, these findings highlight that BACH1 and cMAF cooperate to establish a proinflammatory gene regulatory network in peripheral CD4⁺ T cells in SKO mice, which mirrors the transcriptional signatures of human SLE patients. Loss or inhibition of cMAF is sufficient to ameliorate the SKO phenotype, emphasizing the translational potential of targeting the SATB1-BACH1-cMAF axis in autoimmunity.

## Discussion

Our study identifies a previously unrecognized BACH1-cMAF axis that becomes operative when the genome organizer SATB1 is lost in developing T cells. Using integrated chromatin and transcriptomic profiling, we show that SATB1 normally restricts BACH1 chromatin occupancy in thymocytes; in its absence, BACH1 redistributes toward immune-regulatory loci, enables cMAF binding and together they enforce a proinflammatory transcriptional program. Genetic deletion of *Bach1* in the *Satb1* cKO background reverses the systemic autoimmune-like disease, normalizes cytokine levels, restores tissue architecture and dampens inflammatory gene expression in both the thymus and peripheral CD4⁺ T cells. Consistently, genetic ablation or pharmacologic inhibition of *cMaf* led to a reduction of activated CD4⁺ T cells in SKO mice and additionally the SKO program overlapped human SLE signatures that were diminished in DKO mice, linking the SATB1-BACH1-cMAF axis to lupus-like pathology. These findings provide a mechanistic link between disrupted 3D genome control in the thymus and pathogenic peripheral T-cell states and suggest actionable targets for autoimmune modulation.

The spatiotemporal regulation of transcription factors is essential for T cell lineage commitment and immune homeostasis^9,49–53^. The gene regulatory networks that govern thymocyte development are crucial for the proper function of the adaptive immune response. SATB1 is a T-lineage-enriched genome organizer that shapes promoter-enhancer topology and transcriptional programs during thymocyte maturation^40,54,54^. Prior studies established its high expression at the CD4^+^CD8^+^ stage, its role in 3D genome architecture and its importance in preventing aberrant activation/exhaustion programs in T cells. SATB1 is important for the gene regulatory networks of developing thymocytes through its direct binding in immune-related genes or via the cooperation with major transcription factor complexes such as the NURD complex^55^. These findings highlight a dynamic role between different transcriptional regulators in developing T cells, showcasing their important role in the homeostasis of the adaptive immune response. Conditional *Satb1* loss triggers immune dysregulation and multi-organ inflammation^10^, but the mediators of this breakdown have been incompletely defined. Our data extend this paradigm by showing that SATB1 not only organizes chromatin but also insulates against BACH1 access to immune loci; when SATB1 is absent, BACH1 invades promoter-proximal regions and relocates toward prior SATB1 sites, functionally rewiring thymocyte gene regulation. Strikingly, the inflammatory phenotype is reversed in mice lacking both SATB1 and BACH1 expression, implicating BACH1 as a critical mediator of the pathology.

BACH1 (a CNC-bZIP factor) is best known for heme-sensitive repression of antioxidant and metabolic genes and for forming heterodimers with small MAF proteins. Its roles in adaptive immunity were thought to be indirect, e.g., via effects on antigen-presenting cells or B-cell programs^56^. Direct requirements for BACH1 in T cell chromatin wiring have not been reported. We now demonstrate that BACH1 directly reprograms chromatin in *Satb1*-deficient thymocytes with transcriptional consequences in inflammatory pathways^57^. This reveals a cell-intrinsic BACH1 function in thymocytes and places BACH1 in the causal chain of autoimmunity that follows SATB1 loss.

Loss of SATB1 leads to changes in the chromatin landscape of developing thymocytes and overexpression of molecules that can drive differential gene regulatory networks. The most upregulated gene in the SKO thymocytes is *cMaf*. cMAF is a multifunctional transcription factor in T cells (e.g., Th17/Tfh programs, IL-10 regulation across subsets)^58,24^. cMAF has been identified to regulate the development of follicular helper T cells in the spleen^21,22^ and has also been identified as a crucial factor in the progression of autoimmune neuroinflammation^26,59^. While cMAF has been widely studied, its chromatin recruitment being contingent on BACH1 in thymocytes has not, to our knowledge, been described. Here, cMAF binding is extensive in *Satb1* cKO thymus yet collapses in the *Bach1*-deficient double knockout; at shared sites, occupancy is markedly reduced and SKO-unique cMAF peaks map to immune-regulatory genes that become transcriptionally upregulated. Thus, BACH1 appears to license cMAF access to disease-relevant loci when SATB1 insulation is removed - a cooperative arrangement that explains the emergence of pathogenic thymocyte states and the downstream activation bias of peripheral CD4⁺ T cells.

Our integrated analysis verified that the BACH1/cMAF axis regulates two distinct gene sets in developing thymocytes. The first gene set is activated in the thymus of SKO mice and drives the pathogenic transformation of thymocytes, which become autoreactive, evade T cell selection and ectopically exit towards peripheral lymph nodes and tissues. The second gene set is primed by BACH1/cMAF axis in the thymus, but is upregulated in the peripheral CD4 T cells of SKO spleens, where it promotes proinflammatory cytokine production, neutrophil infiltration, ectopic B cell activation and autoantibody circulation. We propose a model in which SATB1’s architectural function compartmentalizes immune-regulatory chromatin and prevents BACH1 from engaging promoter-proximal and enhancer elements. SATB1 loss (i) increases local accessibility for BACH1, (ii) increases BACH1 promoter proximity and (iii) positions BACH1 near former SATB1 binding sites where it stabilizes cMAF occupancy. The BACH1/cMAF duet then activates cytokine signaling, adhesion and differentiation modules, establishing thymocyte populations with proinflammatory signatures that seed the periphery. This model integrates SATB1’s established roles in 3D genome control with our discovery of BACH1-enabled cMAF recruitment, offering a chromatin-level route from thymic dysregulation to systemic autoimmunity.

The peripheral CD4⁺ T cell programs we observe in *Satb1*-deficient mice mirror SLE-like signatures and align with reports of transcriptional re-wiring in human SLE T cells (including shifts in activation, metabolism, and interferon-linked pathways)^60,61^. While SATB1’s exact expression dynamics in human SLE subsets remain under study, down-tuning of SATB1 activity in pathological contexts and the emergence of cMAF-associated programs have both been implicated in immune dysregulation. Our cross-species comparisons suggest that a SATB1-BACH1-cMAF axis could represent a conserved vulnerability that channels thymic mis-patterning into persistent pathogenic T-cell states.

Because BACH1 is a heme-sensing repressor, it can be pharmacologically modulated. Heme (and hemin) inactivates BACH1 DNA binding, triggers nuclear export and promotes proteasomal degradation^62,63^. Small-molecule BACH1 inhibitors (including HPP-4382 and the selective inhibitor ASP8731) have emerged in oncology and inflammatory contexts and could, in principle, be repurposed to blunt the BACH1/cMAF program in autoimmunity^64,65,66^. On the cMAF side, several groups report small-molecule cMAF inhibitors or repurposed drugs with cMAF-modulatory activity. While preclinical and disease-context work is early, our data nominate cMAF as a tractable node whose activity depends on BACH1 in the *Satb1*-deficient state. A combined strategy - limiting BACH1 availability (e.g., heme mimetics/BACH1 inhibitors) to reduce cMAF recruitment - merits testing in models of T cell-driven autoimmunity.

Conceptually, our work highlights how loss of a genome organizer can expose latent transcriptional circuits that are not inherently pathogenic but become so when redirected to the wrong chromatin neighborhoods. Rather than acting through a single effector, SATB1 appears to buffer multiple transcription factors (here, BACH1 and cMAF) from immune loci. This buffering model may generalize to other contexts where 3D genome stabilizers are perturbed, providing a framework to connect developmental architecture to adult-onset inflammatory disease. While our genetic epistasis places BACH1 upstream of cMAF occupancy, we cannot exclude additional BACH1-independent routes to cMAF recruitment in specialized subsets.

We uncover a SATB1-restrained, BACH1-enabled cMAF program that links thymic chromatin re-patterning to peripheral autoimmune-like pathology. By positioning BACH1 as both a sensor (heme-responsive) and a licensing factor for cMAF, our findings open translational routes to modulate pathogenic T-cell programs upstream of end-organ inflammation. We anticipate that targeting this axis - genetically or pharmacologically - will provide mechanistic leverage to re-establish immune tolerance in SATB1-deficient settings and, potentially, in subsets of human autoimmunity where analogous transcriptional architectures are engaged.

## Data availability

- All data are part of the GEO SuperSeries: GSE308561 (token for review while in private status: avelumkufrozbun)
- Single-cell RNA-seq data of whole thymi have been deposited at GEO at GSE303250 and are publicly available as of the date of publication.
- Single-cell RNA-seq data of whole spleens have been deposited at GEO at GSE303554 and are publicly available as of the date of publication.
- ChIP-seq data have been deposited at GEO at GSE303251 and are publicly available as of the date of publication
- Bulk RNA-seq data have been deposited at GEO at GSE306494 and are publicly available as of the date of publication
- We re-analyzed publicly available ATAC-seq and bulk RNAseq data from WT and SKO thymocytes as well as HIChIP against SATB1 in WT thymocytes (GSE173476)
- We re-analyzed publicly available scRNAseq data from PBMCs of healthy individuals and SLE patients (GSE174188)
- Microscopy data reported in this paper will be shared by the lead contact upon request.
- Any additional information required to reanalyze the data reported in this paper is available from the lead contact upon request.

## Acknowledgments

We would like to thank George Garinis for fruitful discussions. We thank, George Bertsias, Androniki Kretsovali, Joseph Papamatheakis, Eric Pinaud, Iannis Talianidis, Antonis Tatarakis and Lauren Zenewicz for critical reading of the manuscript.

## Author contributions

Conceptualization, C.S., P.T and D.A.P.; methodology, D.A.P., P.T., D.T., E.M., investigation, D.A.P., P.T., D.T., E.M., M.K.; writing - original draft, C.S and D.A.P.; writing - review & editing, C.S., D.A.P., P.T., E.M.; funding acquisition, C.S.; resources, C.S. and K.I.; supervision, C.S.

## Methods

### Mice

All mouse model handlings, breeding, manipulations and sacrifices were approved by the bioethics committee of the IMBB-FORTH and the bioethics committee of the University of Crete in accordance to the rules of the IMBB Animal Facility Committee. The *Satb1^fl^*^/fl^ mice was previously generated and characterized^67^. The *Bach1*^−/-^ (BKO) mouse was previously described. The *Cd4CreMaf*^fl/fl^ mice were kindly provided by Prof. Dr. Birchmeier^68^. The animals used for thymi isolation were 4–8-week-old and for spleen isolation were 90-100 days old. Thymi were harvested and smashed in 10ml 1xPBS using the back of a pestle and homogenized through a 40μm nylon mesh in order to achieve single cell suspension. Cells were centrifuged for 5min at 600g and washed twice with 10ml 1xPBS. These cells were either crosslinked using Para-Formaldehyde (16% methanol free solution, SKU: 15710, Electron Microscopy Sciences) or lysed in order to collect DNA, RNA or protein extracts.

### Protein extract preparation and immunoprecipitation experiments

Freshly isolated thymocytes were lysed in 300ul EBC Lysis Buffer (150mM NaCl, 50mM Tris pH=7.5, 5% Glycerol, 1% Nonidet P-40, 1mM MgCl_2_, 1X Proteinase Inhibitors, 1mM PMSF) for 45 minutes at 4°C with rocking. Samples were centrifuged at 14000rpm for 30 minutes and supernatants were collected in fresh tubes. Protein concentration was measured using Bradford Protein Assay Dye Reagent (Bio-Rad #5000006). For each IP sample 800-1000μg of whole protein extracts were incubated with 5-10μg of antibody for 16-18 hours at 4°C with rocking. 30ul of Magnetic Beads (Pierce™ Protein A/G, Thermo Fischer, cat. No 88803) were washed thrice in EBC Lysis Buffer and incubated with the immunocomplex for 2-4hours at 4°C with rocking. Samples were washed three times with Wash Buffer I (150mM NaCl, 50mM Tris pH=7.5, 5% Glycerol, 0.05% Nonidet P-40, 1mM PMSF) and two times with Wash Buffer II (150mM NaCl, 50mM Tris pH=7.5, 5% Glycerol, 1mM PMSF) for 5 minutes each at 4°C with rocking. Beads were washed twice with TE buffer (10mM Tris pH=8, 1mM EDTA) and resuspended in 40ul SDS-Loading Buffer.

### Nivalenol Treatment

Age-matched WT and SKO mice were treated daily either with 1mg per kilogram of bodyweight of Nivalenol (23282-20-4, Cayman Chemicals) in 4% DMSO or with 4% DMSO for 10 days from day 20 to day 30 after birth. Mice were sacrificed on day 31 and thymi, spleens or lymph nodes were harvested for whole cell extract preparation, RNA extraction and FACS analysis.

### Western Blot

Protein samples in 1X SDS Loading buffer were boiled for 10 minutes at 95°C. Samples were loaded on an 8% SDS-PAGE gel and run for 30 minutes at 90V and for an additional 1.5 hour at 110V. Proteins were transferred in a nitrocellulose membrane at 320mA (constant) for 1.5 hour at 4°C with stirring. Membranes were blocked using 5% Milk/TBS-T for 2 hours and incubated with primary antibodies for 4hours at RT or 16-18hours at 4°C. Membranes were washed three times with TBST for 5 minutes and incubated with 1:2000 secondary antibodies (115-035-146, 111-035-003, 705-035-003, Jackson Laboratories) for 1 hour at RT with rocking. Membranes were incubated with ECL reagent (Pierce Western Blotting Substrate, 32106, Thermo Scientific,) and exposed in the Bio-Rad ChemiDoc Touch Image System Gel Imaging System. Quantitation was performed using ImageJ (Analyze->Gels->Select Lanes->Plot Lanes).

### RNA extraction and cDNA synthesis and RT-qPCR

Freshly isolated thymocytes were resuspended in 1ml Tri Reagent (Sigma-Aldrich, T9424) and incubated for 5m minutes at RT. 250ul of chloroform was added and samples were vortexed for 30seconds and incubated for an additional 5 minutes at RT. Samples were centrifuged at 10.000rpm for 5 minutes in order to collect the aqueous phase. RNA samples were precipitated using 500ul of isopropanol and 5ul of glycogen for 5minutes at RT and tubes were centrifuged at 14000rpm for 20 minutes. RNA pellets were either stored at -80°C in 70% ethanol or resuspended in RNase-free ddH2O for immediate use. For cDNA synthesis, 2μg of total RNA was mixed with 1 µl of oligo(dT)20 (100 µM) and 1ul 10 mM dNTP Mix (10 mM each dATP, dGTP, dCTP and dTTP at neutral pH) and were incubated for 5 minutes at 65°C in a PCR thermocycler. Samples were placed on ice for 1 minute and added 7ul of RT mix (1X RT Buffer, 5mM DTT, 40U RNase Inhibitor (NEB M0314S), 200U RT enzyme (Enzyquest RN012S). Samples were mixed and incubated for 1 hour at 42°C in a PCR thermocycles. RT enzyme was heat inactivated for 15 minutes at 65°C. 50ng of each cDNA sample was used per reaction for 3 technical replicates per biological replicate using SYBR™ Select Master Mix (4472919, Applied Biosystems) and samples were run in a StepOne plus Real-Time PCR (Applied Biosystems). Ct values of each target gene were normalized based on *Hprt* expression levels using the ΔΔCt method. Statistical analysis was performed using one-way analysis of variance (ANOVA) using the foneway function from the *SciPy* package^69^. When the ANOVA indicated significant differences, post-hoc pairwise comparisons were carried out using Tukey’s Honest Significant Difference (HSD) test implemented in pairwise_tukeyhsd from the *statsmodels* package. A significance threshold of p < 0.05 was applied for all analyses.

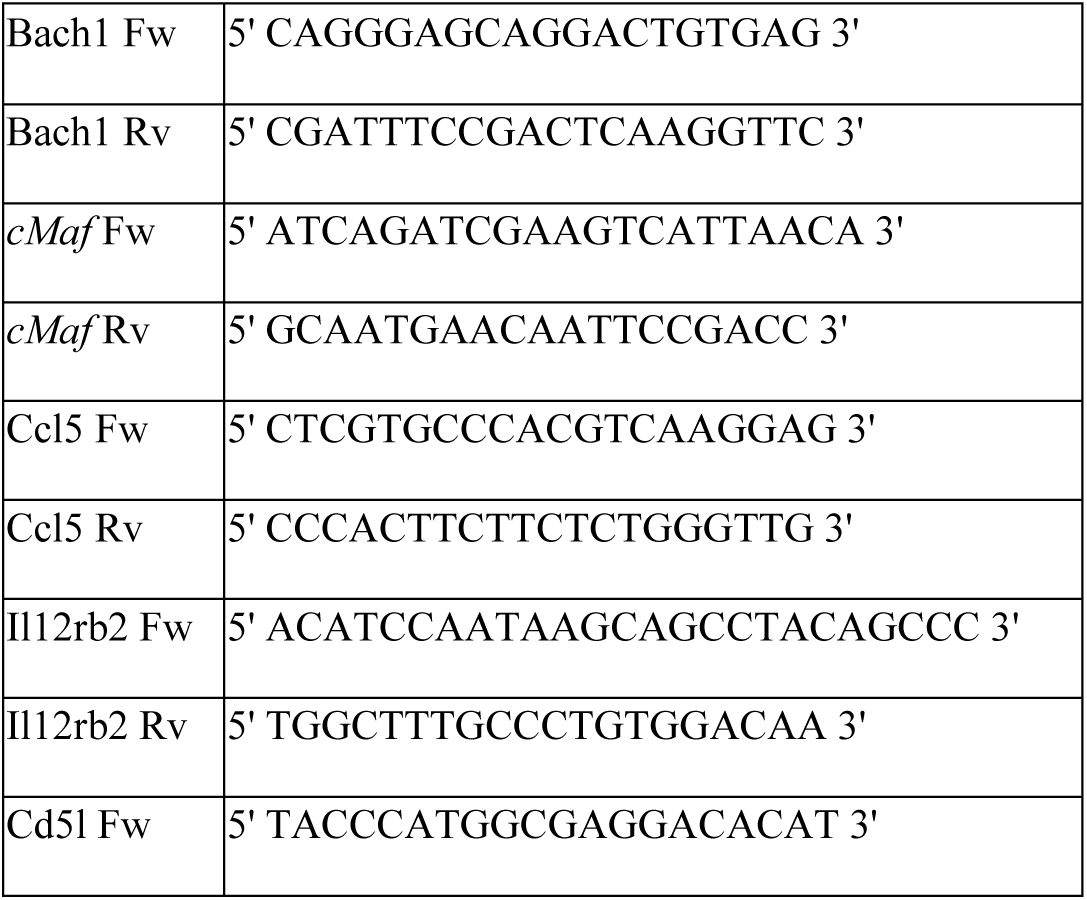
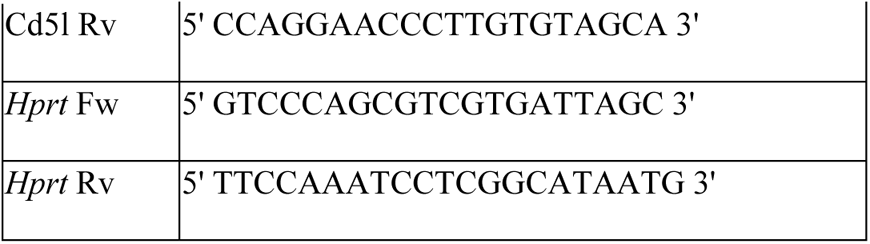

### Histology and Tissue Sectioning

Freshly isolated spleens for 90-100days old mice were isolated and fixed in 4%PFA/1xPBS for 16-18hours at 4°C. Afterwards the tissues were washed with 1xPBS and were dehydrated using 70% Ethanol. The tissues were further dehydrated with two incubations at 90% Ethanol for 30 minutes, three incubations with 100% Ethanol for 30minutes and washed with two incubations in Xylene for 60minutes. Finally, tissues are paraffinized in molten Paraffin for 60mins at 58°C and are embedded in molds. Paraffin blocks are left to harden on a frozen plate and stored at RT in a dark and dry place. Paraffin sections were 5μm thick and placed on positive charged glass slides and let dehydrate at 37°C.

### Flow Cytometry

Staining and FACS analysis were performed as previously^10^. For flow cytometry thymi, spleens or lymph nodes were harvested. Tissues were smashed and single cell suspensions were created using a 40um nylon mesh. Splenocytes were subjected to erythrolysis for 3 seconds in water for red blood cell removal. Cells were counted and 1million was used per sample. All cells were washed once with Staining buffer (1× PBS, 2% FBS, 0.1% NaN3). Thymocytes were incubated with anti-mouse CD4 (APC conjugated, Clone GK1.5, Cat. 100412, Lot. B155935, Biolegend), anti-CD8 (PE conjugated, Clone 53-6.7, Cat. 100708, Lot. B166838, Biolegend) antibodies at 1/100 dilution for 30 minutes at 4°C. Splenocytes and lymph node cells were incubated with anti-CD4 (FITC conjugated, Clone GK1.5, Cat. 100406, Lot. B171662, Biolegend), anti-CD62L (PE conjugated, Clone MEL-14, Cat. 104408, Lot. B169109, Biolegend) and anti-CD44 (APC conjugated, Clone IM7, Cat. 103012, Lot. B17639, Biolegend). Cell population were FSC/SSC gated for the expected size of lymphocytes. Splenocyte populations and lymph node populations were also gated for CD4+ cells. Isotype controls used were Rat IgG2b (FITC conjugated, Clone RTK4530, Cat. 400633, Lot. B265823, Biolegend), Rat IgG2a (PE conjugated, Clone RTK2724, Cat. 400507, Lot. B282895, Biolegend) and Rat IgG2b (APC conjugated, Clone RTK4530, Cat. 400611, Lot. B295805, Biolegend).

### Characterization of cytokine milieu

Cytokines were quantified as previous studies (Zelenka et al. 2022) using the LEGEND-plex Mouse Th Cytokine Panel (Biolegend, 740741). For this study 8 WT, 8 SKO, 8 BKO and 8 DKO female mice of varying age of 1-7 months were used according to the manufacturer’s instructions For each cytokine, pairwise comparisons between genotypes were performed using two-tailed Welch’s t-tests to account for unequal variances.

### Survival Curve

Kaplan–Meier survival analysis was performed to assess survival in wild-type (WT, n = 14), single knockout (SKO, n = 18), and double knockout (DKO, n = 17) mice. Animals were monitored daily from birth, and survival was recorded as the number of days lived until natural death or euthanasia for humane endpoints. Survival curves were generated using the Kaplan–Meier method, with censored animals indicated on the plots. Statistical comparisons between groups were performed using the log-rank (Mantel–Cox) test.

### Autoantibody Detection

Paraffin sections from WT pancreas were deparaffinized for 30 minutes at 55°C and were washed twice with Xylene for 3 minutes. Slides were incubated in 100%/100%/95%/70% Ethanol for 3 minutes. Antigen retrieval was performed using 10mM sodium citrate pH 6 + 0.05% Tween-20 and slides were incubated twice for5 minutes at 80°C. Slides were let to cool down and were incubated in NGS blocking buffer (5% Normal Goat Serum in TBST) for 30 minutes at RT. Sera from WT, SKO and DKO mice were diluted 1:10 in NGS blocking buffer. Slides were incubated with sera for O/N in a humidified chamber at 4°C. Slides were washed twice with TBST for 5 minutes at RT and were incubated with Alexa 594 Goat anti-mouse IgG (H+L) secondary antibody (Cat # A-11032, Invitrogen) in NGS Blocking Buffer for 60 minutes at RT. Slides were washed three times with TBST for 5 minutes and were incubated with 1μM DAPI(Cat# P-36931, Invitrogen) in NGS Blocking Buffer for 10 minutes at RT. Slides were washed three times with TBST and mounted with Mowiol (81381 Sigma-Aldrich). Imaging was performed in a Leica SP8 confocal microscope.

### Hematoxylin and Eosin Staining

Paraffin sections were deparaffinized for 30-60 minutes at 55°C. Slides were washed three times with Xylene for 3min at RT and washed twice with 100% Ethanol and once with 95% ethanol for 3 minutes at RT. Slides were rinsed in ddH2O for 1 minute and stained with Hematoxylin for 2.5 minutes at RT. Slides were washed with ddH2O for 3 minutes and were differentiated in 5% acetic acid/70% ethanol for 45 seconds. Slides were washed once with ddH2O for 1 minute and incubated in Bluing Buffer (0.2% w/v KHC03, 2% w/v MgSO4) for 30 seconds. Slides were washed in ddH2O for 2 minutes, 95% Ethanol for 45 seconds and stained with Eosin for 30-45 seconds. Eosin was washed away with 3 washes at 100% ethanol for 2 minutes and 2 washes with xylene for 3 minutes. Mounting was performed using Estellan mounting medium (1.07960, Sigma-Aldrich).

### Thymocytes Immunofluorescence

Glass coverslips were coated using 0.1mg/ml Poly-D-Lysine. Freshly isolated thymocytes were coated at a density of 500-800 thousand cells per coverslip and centrifuged at 500rpm for 3 minutes and washed with 1xPBS. Cells were crosslinked with 4% PFA/1xPBS for 10min on ice and washed 3 times with 1xPBS. Cells were permeabilized with 0.5% Triton-X/1xPBS for 5min on ice and washed 3 times with 1xPBS. Cells were incubated with Blocking Buffer (0.4% acetylated BSA, 4xSSC, 0.1% Tween-20) for 30minutes at RT in humidified chamber. Afterwards cells were incubated with primary antibodies in Detection Buffer (0.1% acetylated BSA, 4xSSC, 0.1% Tween-20) for 1 hour at RT in humidified chamber and washed 3 times with Washing Buffer (4xSSC, 0.1% Tween-20). All Secondary antibodies were used in final concentration of 1:250 in Detection Buffer for 45 minutes at RT in humidified chamber. Cells were washed 2 times with Washing Buffer incubated with 1:2000 DAPI (stock 5mg/ml), washed once more with washing buffer and mounted using 10ul Mowiol mounting medium. Nuclear versus cytoplasmic signal quantitation of cMAF immunofluorescence experiments was performed using Cyt/Nuc ImageJ macro^70^. Briefly the individual channels of an image are split and the threshold is set based on the saturation of the DAPI (Cat# P-36931, Invitrogen) staining in order to define the nuclear region versus the cytoplasmic. The nuclear masks were used to define cytoplasmic regions by exclusion. Two-way ANOVA was performed to evaluate the effects of cellular compartment (nuclear vs. cytoplasmic) and experimental condition on fluorescence intensity measurements. The analysis was conducted using GraphPad Prism 8 with compartment as the row factor and condition as the column factor. Interaction effects between factors were also assessed. Post-hoc comparisons of means were carried out where appropriate, and statistical significance was defined as *p* < 0.05.

### Mitotracker staining of coverslip-coated thymocytes

Freshly-coated thymocytes were washed once with 1xPBS and incubated with 500nM Mitotracker (Invitrogen Cat#M7514) for 15 minutes at 37°C. Cells were washed three times with 1xPBS and were fixed with 4% PFA/1xPBS for 10 minutes on ice. Cells were washed two times with 1xPBS, stained with 1:1000 DAPI (5mg/ml, Cat# P-36931, Invitrogen) for 10 minutes at RT and finally washed once more with 1xPBS. Coverslips were mounted on glass slides using Mowiol mounting medium. Images were analyzed using ImageJ (Mean Fluorescent Intensity).

### Spleen paraffin sections

Paraffin sections were incubated for 30-60 minutes at 55°C. Slides were washed three time with xylene for 10 minutes at RT and were rehydrated with consecutive washes in 100%/95%/90%/80%/70% ethanol for 3 minutes. Slides were incubated twice in prewarmed Antigen Retrieval Buffer (10mM sodium citrate pH=6 /0.05% Tween-20) at 100°C for 5 minutes. Slides were left to cool down and washed twice with 1xPBS for 5 minutes at RT with shaking. Tissue sections were permeabilized with 0.3%Triton/ 1xPBS for 30 minutes at RT with shaking. Afterwards sections were blocked with 5%NGS/ 0.3%Triton/ 1xPBS for 1 hour at RT in humidified chamber. Primary antibodies were incubated in 1%NGS/ 0.3%Triton/1xPBS for O/N at 4°C in humidified chamber. Slides were washed three times with 0.3%Triton/1xPBS for 10 minutes at RT with rocking and incubated with secondary antibodies in 1%NGS/ 0.3%Triton/ 1xPBS for 1 hour at RT in humidified chamber. Slides were washed twice with 0.3%Triton/1xPBS for 10 min at RT with shaking, stained with 1:1000 DAPI (Cat# P-36931, Invitrogen) for 10minutes at RT and after a final wash they were mounted with glass coverslips with Mowiol.

### Chromatin immunoprecipitation (ChIP)

Freshly isolated thymocytes in 10ml 1xPBS were crosslinked with 1ml Fixation Buffer (11% Formaldehyde, 100 mM NaCl, 1 mM EDTA, 0.5 mM EGTA, 50 mM Hepes pH 8.0) for 10 minutes at RT rotating. PFA was quenched with 0.2M Glycine for 5 minutes at RT, rotating and samples were wshed two times with 10ml 1xPBS and centrifuged at 1000g for 5 minutes at 4°C. Crosslinked cell pellets were either flash-frozen in liquid nitrogen and stored at -80°C or were proceeded for ChIP. Each ChIP sample contained 15million thymocytes and were lysed with 60ul ChIP Lysis Buffer (1% SDS, 50 mM Tris (pH 8), 20 mM EDTA, 1X PIC=protease and phosphatase inhibitors) for 20 minutes at RT. SDS was then diluted with the addition of 540ul TE Buffer + 1XPIC and placed on ice. Samples were sonicated for 3-4 minutes (30sec ON/OFF cycle, 40% amplitude) on a Labsonic-M Tip Sonicator. Samples were centrifuged at 14000g for 30 minutes at 4°C and the supernatant was collected. Samples were dialyzed for three consecutive 2-hour incubations in Dialysis Buffer (10 mM Tris pH 8.0, 1 mM EDTA, 150 mM NaCl, 0.1% Na-Deoxycholate, 1mM PMSF). In parallel 30-40ul of magnetic beads per sample were washed three times with BSA/PBS Buffer (0.1% BSA/1xPBS) and were incubated with 8-10ug of desired antibody for 6 hours at 4°C. Samples were collected and incubated with 1% Triton-X for 10 minutes at 37°C. Each sample was added an equal volume of 2X ChIP Binding Buffer (20 mM Tris pH 8.0, 2 mM EDTA, 300 mM NaCl, 0.2% Na-Deoxycholate, 2X PIC). Bead-Ab complexes were washed three times with BSA/PBS Buffer and combined with the chromatin samples for 16hours at 4°C rotating. Beads were washed seven times with RIPA Buffer (50 mM Hepes (pH 8.0), 1% NP-40, 0.70% Na-Deoxycholate, 0.5 M LiCl, 1X PIC, 1 mM EDTA), two times with TE and were resuspended in 125ul ChIP Elution Buffer (10 mM Tris-HCl (pH 8.0), 5 mM EDTA, 300 mM NaCl and 1% SDS). Samples were de-crosslinked for 18 hours at 65°C and treated with 100ug RNase A for 30 minutes at 37°C and 80ug Proteinase K for 3 hours at 55°C. Samples were cleaned either with Phenol:Chloroform extraction or with the Zymo Research ChIP DNA Clean and Concentrator kit (D5205) and resuspended in ddH2O.

| Antibody | Supplier | Cat. No | lot |
| --- | --- | --- | --- |
| anti- SATB1 | custom (Davids Biotechnologies) | custom | - |
| anti- BACH1 | custom (Davids Biotechnologies) | custom | - |
| anti-cMAF | Santa Cruz | 293420 | 1524 |
| anti-cMAF | Bethyl | A300-613A | 2 |
| anti-H3K4me1 | Abcam | ab8896 | GR127871-2 |

### DNA Library Preparation for NGS sequencing

20-50ng of ChIP material was used per sample. DNA was mixed with 1.25-2.5ul Tn5 enzyme (Illumina Tagment DNA Enzyme and Buffer Small Kit, 20034197) 2X TD Buffer (20 mM Tris pH 7.5, 10 mM MgCl2, 20% Dimethylformamide) and ddH2O to a final volume of 50ul. Samples were incubated for 10 minutes at 55°C with time-to-time pipetting. Reaction was stopped with 10ul Strip Buffer (0.6% SDS, 60mM Tris pH=8.0, 300mM EDTA) and incubated for 5 minutes at RT. Samples were cleaned using Zymo Research ChIP DNA Clean and Concentrator kit (D5205) and resuspended in 24ul ddH2O. The 1/10 of this material was used in a pilot qPCR to determine the number of PCR cycles each sample. Each sample was used for a PCR reaction using 1.5ul of each forward or reverse specific indexed primers (Nextera DNA Sample Preparation Index Kit, Illumina, FC-121-1011), 25ul 2X Phusion Master Mix and ddH2O to a final volume of 50ul. The PCR reaction was performed following the program 72 °C for 5minutes and repeated cycles of 98 °C for 15 seconds, 63 °C for 35 seconds, 72 °C for 1minute.

### Data Processing

Raw reads were trimmed (adapter trimming, low quality trimming of phred<20, <20bp nucleotide removal) using <u>trim-galore</u> and mapped to mm10 with the use of hisat2^71^ at default parameters. Samples were sorted, indexed and filtered using samtools^72^. Multi-mapped reads and duplicate reads were also discarded using samtools and peaks were called using macs2^73^ (input normalization, nomodel, --extsize 147). Genome coverage was generated using deepTools2^74^ (normalization based on RPKM, bin=50). BACH1 and cMAF BigWig sample scores were plotted to genomic regions of interest (BACH1 peaks, cMAF peaks, thymocyte enhancers, SATB1 peaks) using the computeMatrix and plotProfile functions. Overlapping peaks between samples were generated using bedtools^75^. Differential binding or deposition of a factor or mark on the same genomic regions was found with the use of diffbind^76^ and DESeq2^77^ while GO term enrichment of these differential peaks was assessed using ClusterProfiler^78,79^.

### BACH1 peak length estimation and distance from SATB1 peaks

BACH1 peak files were reformatted into BED format with four columns (chromosome, start, end, and peak length). For each condition, peak lengths were calculated as the difference between the end and start coordinates. Length distributions of BACH1 peaks in WT and SKO were compared using statistical testing (Student’s *t*-test or Wilcoxon rank-sum test, depending on normality). To evaluate the spatial proximity of BACH1 and SATB1 binding sites in WT and SKO thymocytes, the genomic distance between BACH1 and SATB1 peaks was calculated using BEDTools’ closest function, identifying the nearest SATB1 peak for each BACH1 peak. SATB1 peaks were retrieved from previous studies^10^ (GSE173476). Resulting distance distributions were visualized and statistically compared as above.

### Bulk-RNAseq sequencing

Freshly isolated thymocytes form 4–6-week-old C57B/6 and BACH1^−/-^ mice were washed twice with 1X PBS and resuspended in 1ml TRI Reagent (AM9738, Invitrogen). 250ul of chloroform were added per 1ml of TRI Reagent and the aqueous phase was collected in a tube. Samples were precipitated with equal volume of isopropanol by incubation for 20 minutes on ice and centrifugation at 12000g for 20 minutes. RNA pellets were washed twice with 75% Ethanol and resuspended in ddH2O. Samples were treated with 20 Units of DNase for 20 minutes at 37°C and were purified using RNeasy Mini Kit (Qiagen, 74104). Libraries and sequencing were performed in the Greece Genome Center in the Bioacademy of Athens.

### Bulk-RNAseq data processing

Raw reads were mapped to the mm10 reference genome using Bowtie2^80^. Multi-mapped reads and duplicate reads were discarded using samtools. Feature counts^81^ were used to summarize the reads of each transcript and gene and differential expression analysis was performed using DESeq2^77^. Gene ontology analysis was performed using Cluster Profiler^78,79^ and volcano plots were generated using EnhandedVolcano^82^.

### ATAC-seq data processing

Accessibility regions from WT and SKO thymocytes were utilized from previous publicly available data of the lab^10^.

### Single-cell RNAseq analysis

Freshly isolated thymi or spleens (2 biological replicates per genotype) were harvested and washed three times with 1xPBS and shipped in cryopreservation buffer. Samples were sent to Singleron Biotechnologies (Cologne, Germany) for tissue dissociation, single cell suspension, single cell capture, cDNA preparation from single cells, library preparation and NGS sequencing. We aimed for 10.000 cells per sample for scRNAseq and 5000 cells for scTCRseq. Each cell was destined for 35.000 reads. Raw reads were analyzed with the Celescope pipeline (https://github.com/singleron-RD/CeleScope). Doublets were removed with the use of DoubletFinder^83^. Cells with >5% mitochondrial counts were removed and samples were processed using the standard pipeline of Seurat^84^ (NormalizeData, FindVariableFeatures, ScaleData,RunPCA, IntegrateLayers, JoinLayers, FindNeighbours, FindClusters, RunUMAP). Cluster markers were found using FindMarkers (min.pct=0.25) and volcano plots were generated with the R package EnhnancedVolcano (fc>0.5, pvalue<0.05). Gene enrichment analysis (fc>2, pvalue<0.05) and dotplots was performed with ClusterProfiler. Pseudo-bulking counts were analyzed with DESeq2^83^. Cell identities were annotated using the SingleR package (reference from ImmGen Consortium, sorted bulk RNAseq, label.fine)^85^. Cell–cell communication analysis was performed using the CellChat R package using the mouse ligand-receptor interaction CellChatDB.mouse database. First, overexpressed signaling genes were identified, and the probability of intercellular communication was computed at the level of individual ligand–receptor interactions as well as signaling pathways (identifyOverExpressedGenes, computeCommunProb, computeCommunProbPathway, aggregateNet. Chord diagrams were generated to visualize outgoing signaling patterns of T cells and B cells in each condition. Cells were either grouped based on clusters (thymus) or based on SingleR identity (spleen). Bach1/cMaf expressing cells were subset based on expression, retaining only those with log-normalized expression values greater than 0.5.

scRNAseq data from thymus were used to perform single-sample Gene Set Enrichment Analysis (ssGSEA) using the GSVA R package^86^ and mouse hallmark gene sets were obtained from the msigdbr database. ssGSEA scores were calculated with a Gaussian kernel and absolute ranking of expression values. The resulting pathway activity scores were incorporated into the Seurat object as a new assay, scaled, and visualized using heatmaps, feature plots, and violin plots to compare pathway activity across genotypes (WT, SKO, DKO).

### scRNAseq analysis from SLE patients

To identify the correlation of expression of *SATB1*, *BACH1* and *cMAF* in autoimmune-disease patients we retrieved the publicly available data of healthy and SLE patient scRNAseq from Perez et al. 2022 (GSE174188). The data were analyzed using scanpy^87^ and cells were sorted (.isin) in order to subset them based on solely *SATB1* expression (removing *BACH1*/*cMAF* expressing cells) or solely on *BACH1* and/or *cMAF* expression (removing *SATB1* expressing cells). Cell populations and percentages were plotted using sc.pl.umap, sc.pl.dotplot and matplotlib^88^. The T cells from the SLE patients were subset and the most upregulated genes were used as an SLE signature. These genes were added in our scRNAseq data from spleens using the AddModuleScore command. For each cell this function computes the average expression levels of the SLE signature minus the average expression levels of highly expressed control gene sets. The module score of each cell is stored in a new metadata column and data were visualized using the FeaturePlot command and SLE_signature as a feature.

### *In silico* modeling of SATB1/BACH1/cMAF using AlphaFold

Structure modelling of BACH1 and cMAF was performed using Alphafold 2.3^89^ colab python script and Alphafold3 Server. The amino-acid sequence of BACH1 and cMAF were matched to publicly available databases (uniref90, mgnify, uniport, smallbfd) and Alphafold was run for 20 recycles in order to ensure the predicted structure confidence. Alphafold predicted structures were visualized and plotted using ChimeraX^89^.

