## Supplementary Figures for "Genome organizer SATB1 restrains the BACH1-cMAF axis that drives autoimmunity"

### Extended Data - Figure titles and legends

#### Extended Data Fig. 1

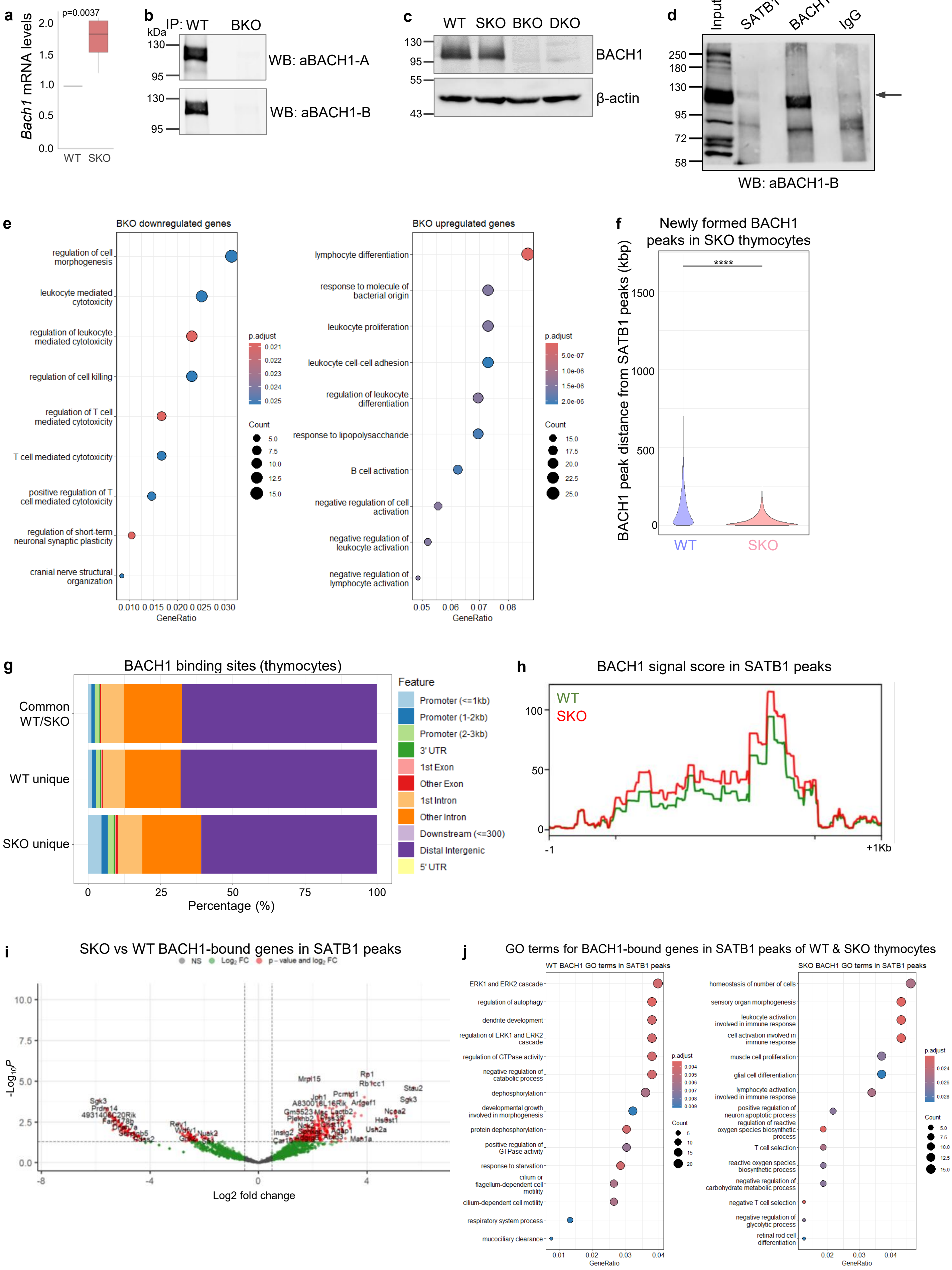

**Extended Data Fig. 1**

**Validation of BACH1 expression and binding in thymocytes**

**a.** RT-qPCR for the relative Bach1 mRNA levels in WT and SKO thymocytes normalized to Hprt1 levels (Wilcoxon rank-sum test, n=3). **b.** Western blot analysis of BACH1 immunoprecipitated from whole-cell thymocyte extracts of WT and *Bach1*$^{-/-}$(BKO) mice (n=4). Antibody A targets the N-terminus, while antibody B targets the C terminus of BACH1. **c.** Western blot analysis for the BACH1 protein expression levels in whole-cell protein extracts from C57BL/6 (WT), Cd4Cre-*Satb1*$^{fl/fl}$ (SKO), *Bach1*$^{-/-}$ (BKO) and double *Cd4*$^{Cre}$-*Satb1*$^{fl/fl}$/*Bach1*$^{-/-}$ (DKO) thymocytes (n=5). **d.** Western blot analysis for BACH1 co-immunoprecipitated with either SATB1, BACH1 or IgG from whole cell thymocyte extracts (n=2). **e.** GO terms from differential gene expression analysis of BKO vs WT thymocytes from bulk RNAseq data (Wald test, FDR-adjusted pvalue, padj<0.05). **f.** Proximity analysis of SATB1 and BACH1 ChIPseq binding sites (peaks) in WT and SKO thymocytes. For SATB1 the HiChIP SATB1 binding sites have been utilized (Wilcoxon rank sum test, p-value < 2.2e-16). **g.** Feature annotation for BACH1 ChIPseq binding sites in WT/SKO common or WT- and SKO-unique BACH1 peaks. **h.** Summary Plot of WT/SKO BACH1 ChIPseq binding profile in SATB1 peaks (summit 5Kb +-1Kb). **i.** Volcano Plot of the differential binding analysis for BACH1 on SATB1 peaks in SKO/WT thymocytes (Exact test, pvalue<0.05, fc>0.5). **j.** Gene ontology of the differentially bound genes by BACH1 in SATB1 peaks of WT and SKO thymocytes (Exact test, FDR-adjusted pvalue, padj<0.05).

#### Extended Data Fig. 2

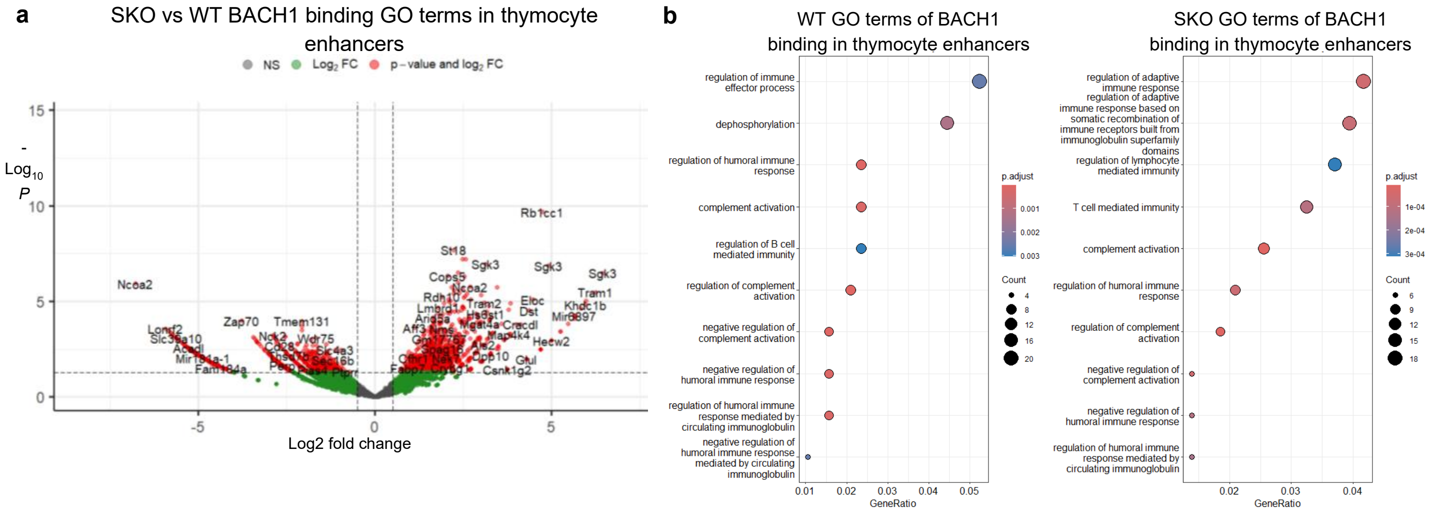


#### Extended Data Fig. 2

##### BACH1 binding at open chromatin and enhancers

**a.** Volcano Plot for the differential binding of BACH1 in thymus enhancer regions in SKO vs WT thymocytes (Exact test, pvalue<0.05, fc>0.5). **b.** GO terms of genes near thymocyte enhancers bound by BACH1 in SKO vs WT thymocytes (Hypergeometric test, FDR-adjusted pvalue, padj<0.05). **c.** Gating strategy for thymocyte FACS analysis of CD4 and CD8 positive populations. **d.** Gating strategy for spleen or lymph nodes FACS analysis of CD4 positive cells expressing CD62L and CD44 antigens.

### Extended Data Fig. 3

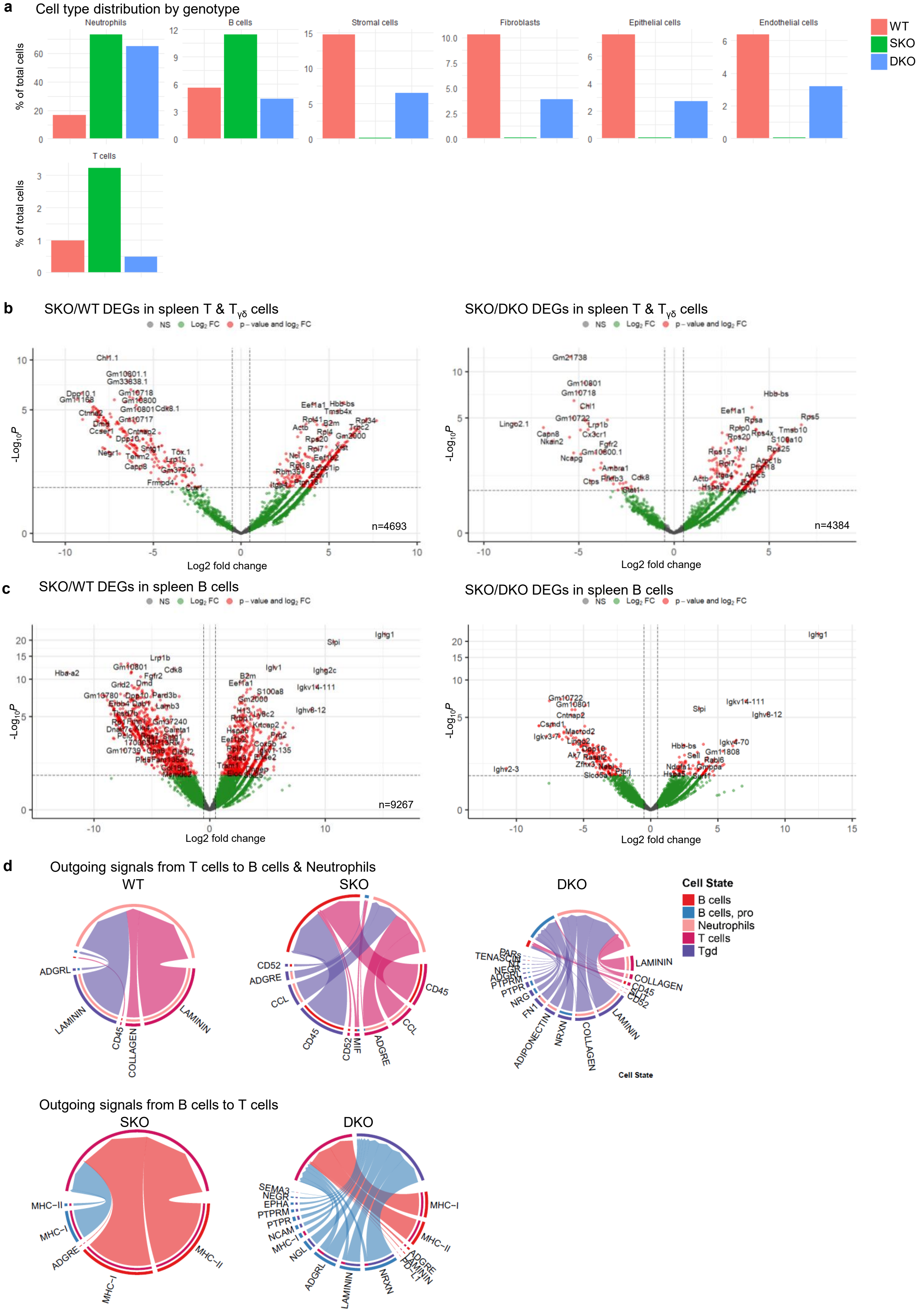

**Extended Data Fig. 3**

**Expanded analysis of splenic immune populations**

**a.** Cell type distribution (percentage of cells) in WT, SKO and DKO spleens in scRNAseq. **b.** Volcano Plot of differential gene expression analysis between SKO/WT and SKO/DKO for T and Tγδ cells of spleen scRNAseq data (Wald test, pvalue<0.05, fc>0.5). **c.** Volcano Plot of differential gene expression analysis between SKO/WT and SKO/DKO B cells in spleen scRNAseq data (Wald test, pvalue<0.05, fc>0.5). **d.** Cellchat analysis of all the outgoing signals from T cells to B cells and neutrophils in WT, SKO and DKO mice, from spleen scRNAseq data (upper panel). Cellchat analysis of all the outgoing signals from B cells to T cells in SKO and DKO mice from spleen scRNAseq data (lower panel).

#### Extended Data Fig. 4

**a** Cell type annotation in thymus scRNAseq

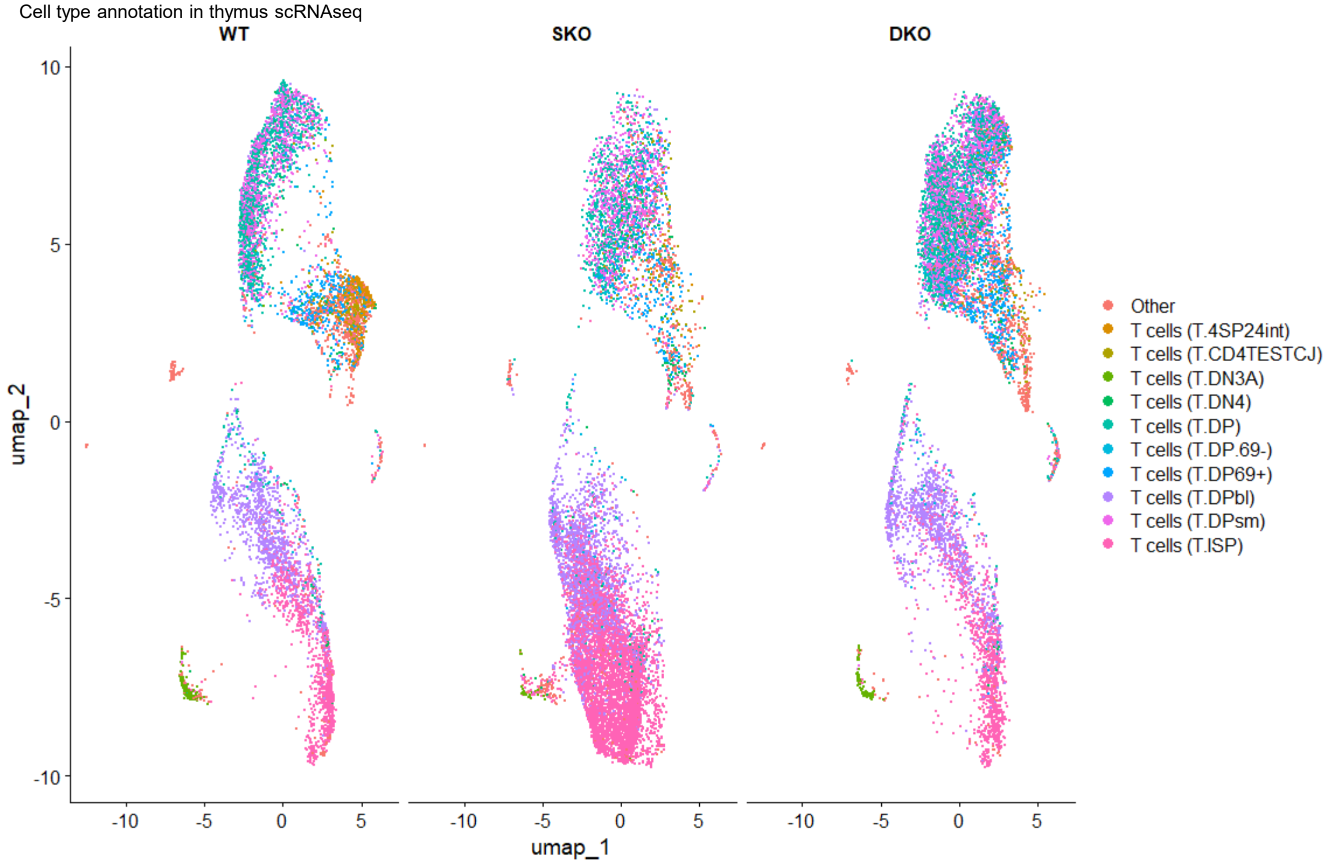


**b** Cell type percentages in thymus scRNAseq

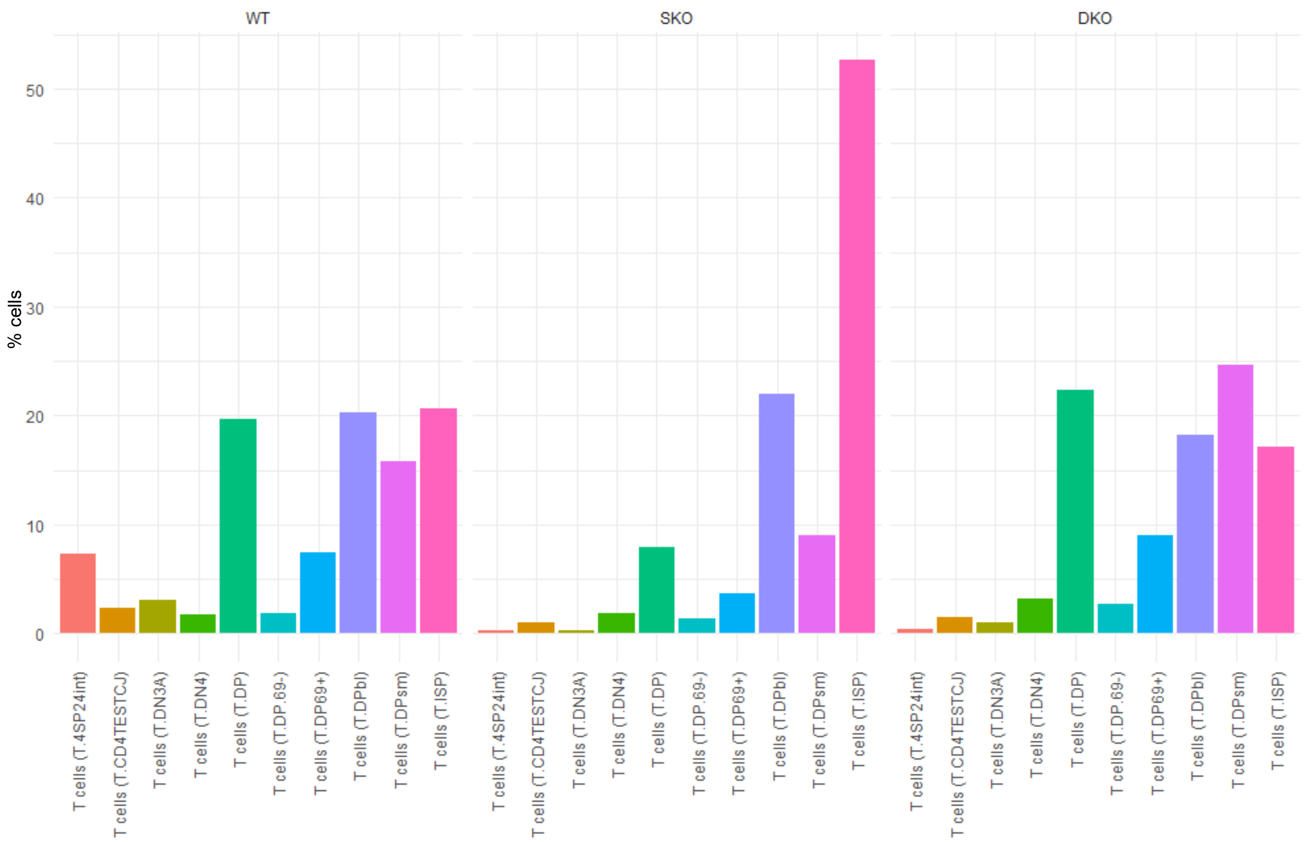

**Extended Data Fig. 4**

**Cell type annotation in thymus scRNAseq**

**a.** Annotation of top ten cell types in thymus scRNAseq from WT, SKO and DKO mice. T cell types: 4SP2int: CD4 single positive 2 intermediate, CD4TESTCJ: CD4 T effector / selection cluster J, DN3A: double negative stage 3A, DN4: double negative stage 4, DP: double positive $CD4^{+}CD8^{+}$, DP69-: double positive ($CD4^{+}CD8^{+}$) $CD69^{-}$, $DP69^{+}$: double positive ($CD4^{+}CD8^{+}$) $CD69^{+}$, DPbl: double positive ($CD4^{+}CD8^{+}$) blast, DPsm: double positive ($CD4^{+}CD8^{+}$) small, ISP: immature single positive. **b.** Percentage of cells coexpressing Bach1 and cMaf in WT, SKO and DKO thymocytes.

#### Extended Data Fig. 5

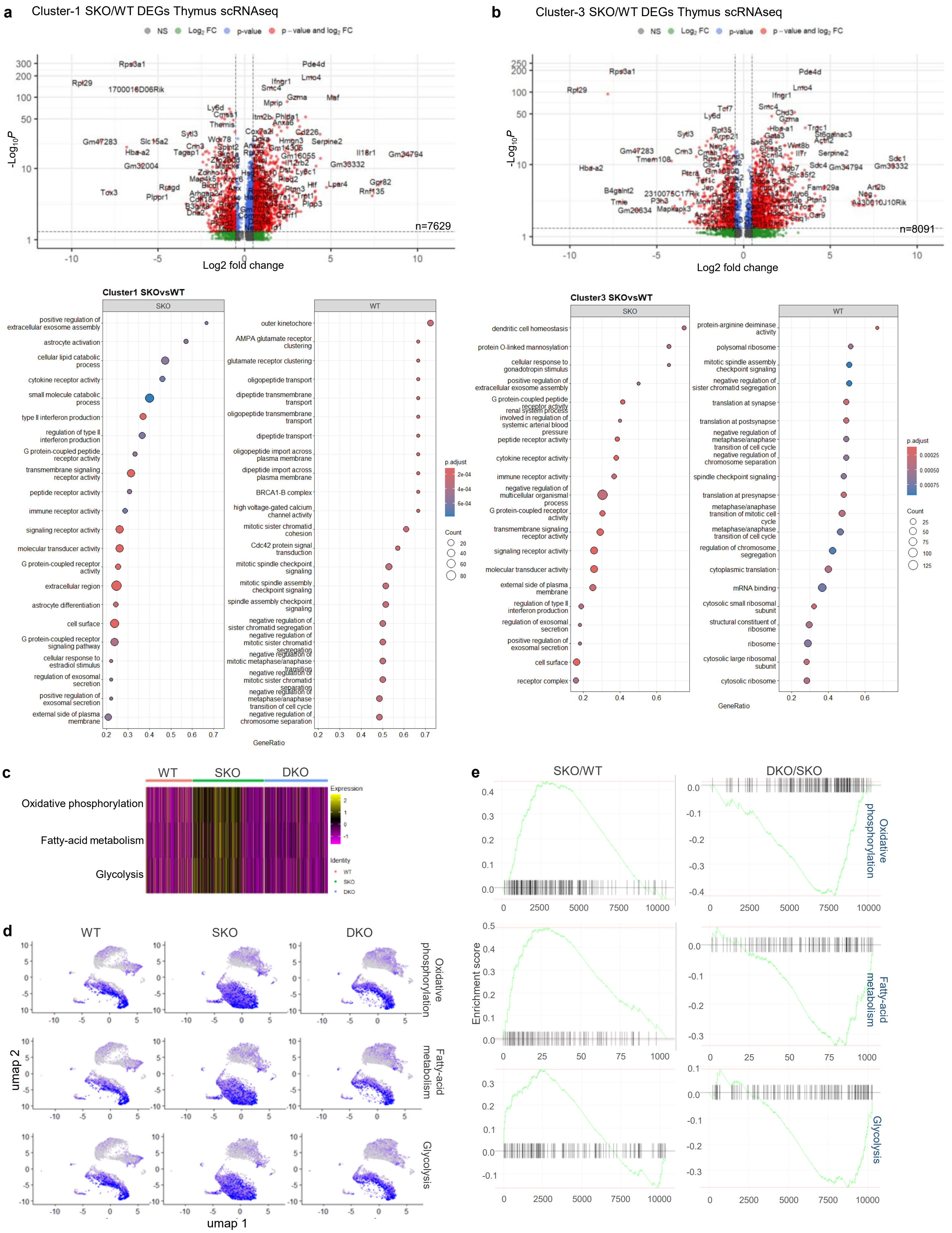

**Extended Data Fig. 5**

**Differential expression analysis of thymocyte population in SKO thymi**

**a.** Volcano plot and gene ontology analysis of the differentially expressed genes of cluster 1 in SKO/WT thymocytes from whole thymi scRNAseq data (Wilcoxon rank sum test, pvalue<0.05, fc>0.5) (Hypergeometric test, FDR-adjusted pvalue, padj<0.05). **b.** Volcano plot and gene ontology analysis of the differentially expressed genes of cluster 3 in SKO/WT thymocytes from whole thymi scRNAseq data (Wilcoxon rank sum test, pvalue<0.05, fc>0.5) (Hypergeometric test, FDR-adjusted pvalue, padj<0.05) (Linked to Fig4A). **c.** GSEA analysis of whole thymi scRNAseq data in WT, SKO and DKO mice. **d.** UMAP visualization of the oxidative phosphorylation, fatty acid metabolism and glycolysis score in scRNAseq data from WT, SKO and DKO thymi. **e.** Enrichment plot from GSEA in whole thymi scRNAseq data in SKO/WT or DKO/SKO thymocytes for oxidative phosphorylation, fatty acid metabolism and glycolysis.

### Extended Data Fig. 6

**a**

| Gene | Locus | WT FPKM | SKO FPKM | Log2 Fold change | q_value |
|---|---|---|---|---|---|
| *Maf* | 8:115682941-115707794 | 0.34 | 64.77 | 7.58 | 0.00111 |
| *Sdc1* | 12:8771322-8793715 | 0.64 | 61.24 | 6.59 | 0.00111 |
| *Actn2* | 13:12269159-12340760 | 0.24 | 17.12 | 6.13 | 0.00111 |
| *Ankrd35* | 3:96670130-96691033 | 0.06 | 4.22 | 6.07 | 0.00111 |
| *Hlf* | 11:90336535-90390895 | 0.11 | 7.26 | 6.06 | 0.00111 |

**b**

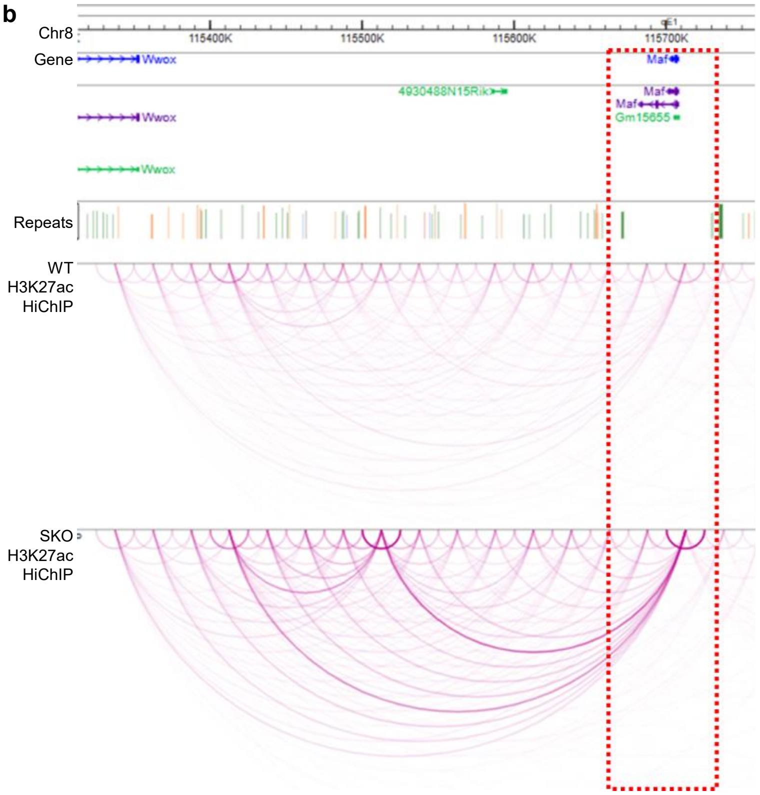


**c**

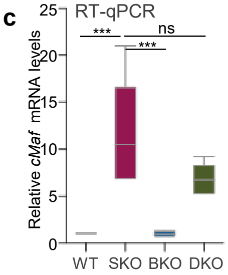


**d**

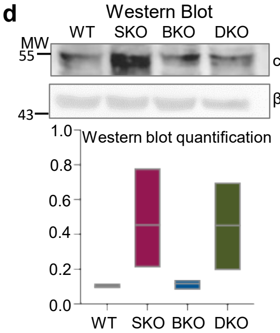


**e**

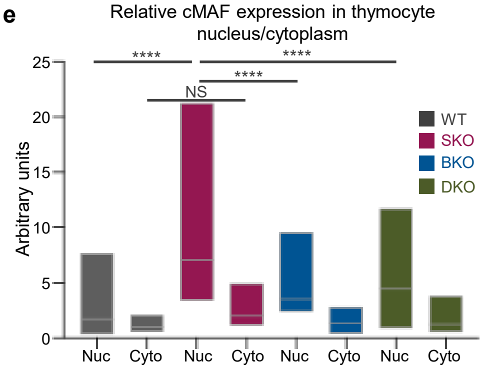


**f**

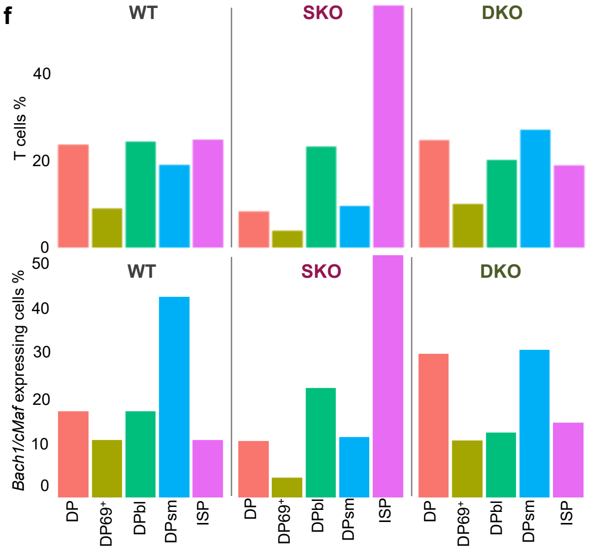


**g**

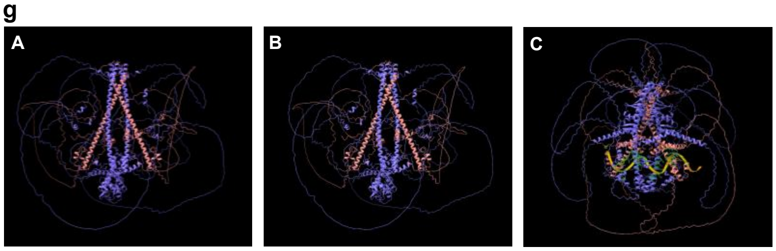


**h**

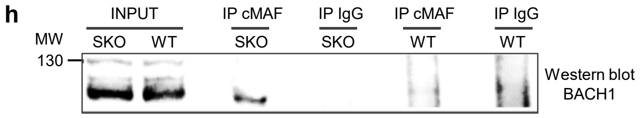


**i**

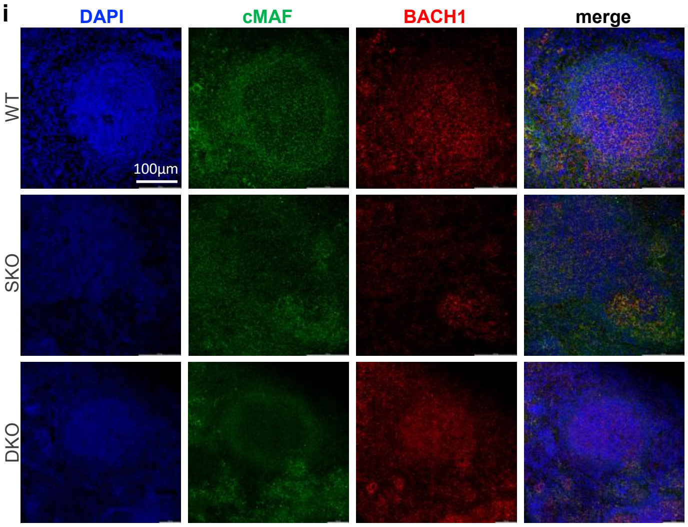

**Extended Data Fig. 6**

**cMAF expression and interaction with BACH1**

**a.** Table of the top 5 upregulated genes in SKO/WT thymocytes from bulk RNAseq analysis. **b.** Genome browser tracks of the *cMaf* locus from H3K27ac-associated chromatin loops (HiChIP) in WT and SKO thymocytes. **c.** RT-qPCR, of *cMaf* mRNA levels in WT, BKO, SKO and DKO thymocytes normalized to Hprt1 levels (two-way ANOVA, mean ± SEM). **d.** Western blot for cMAF levels from thymocyte protein extracts from WT/BKO/SKO/DKO and densitometric quantitation. **e.** Quantitation of the relative protein expression levels of cMAF (based on immunofluorescence) in the nucleus and the cytoplasm of WT, BKO, SKO and DKO thymocytes (n=5, data represent the mean ± SEM). A two-way ANOVA was used to assess the effects of compartment (nuclear vs. cytoplasmic) and treatment condition (four groups). (Post-hoc multiple comparisons were performed with Tukey's test). **f.** Cell identity of the *cMaf*-expressing cells in thymus scRNAseq. This analysis highlights a distinct *cMaf* expressing population blocked in the T ISP stage before DP T cell stage in SKO thymi (chi-squared tests for proportions, all cells T ISP: WTvsSKO: p-value < 2.2e-16, SKO/DKO: p-value < 2.2e-16, WT/DKO: p-value = 1.003e-05, *Bach1*/*cMaf* expressors T ISP: WTvsSKO: p-value = 7.682e-05, SKO/DKO: p-value < 2.2e-16S, WT/DKO: p-value = 0.3476). T cell types: DP: double positive $CD4^+CD8^+$, $DP69^+$: double positive (CD4+CD8+) CD69+, DPbl: double positive (CD4+CD8+) blast, DPsm: double positive ($CD4^+CD8^+$) small, ISP: immature single positive. **g.** AplhaFold simulations of BACH1 and cMAF binding. A and B are image and movie of a tetramer comprised of 2 BACH1 and 2 cMAF molecules. C is the movie of a tetramer comprised of 2 BACH1 and 2 cMAF molecules and an Rhs6 DNA element of the TH2 LCR. **h.** Coimmunoprecipitation experiments against BACH1 and cMAF from murine whole-cell thymocyte protein extracts (input is 10% of whole thymocyte extract immunoprecipitated). **i.** Immunofluorescence against BACH1 and cMAF in spleen follicles of WT, SKO and DKO spleen sections (n=3).

**Extended Data Fig. 7**

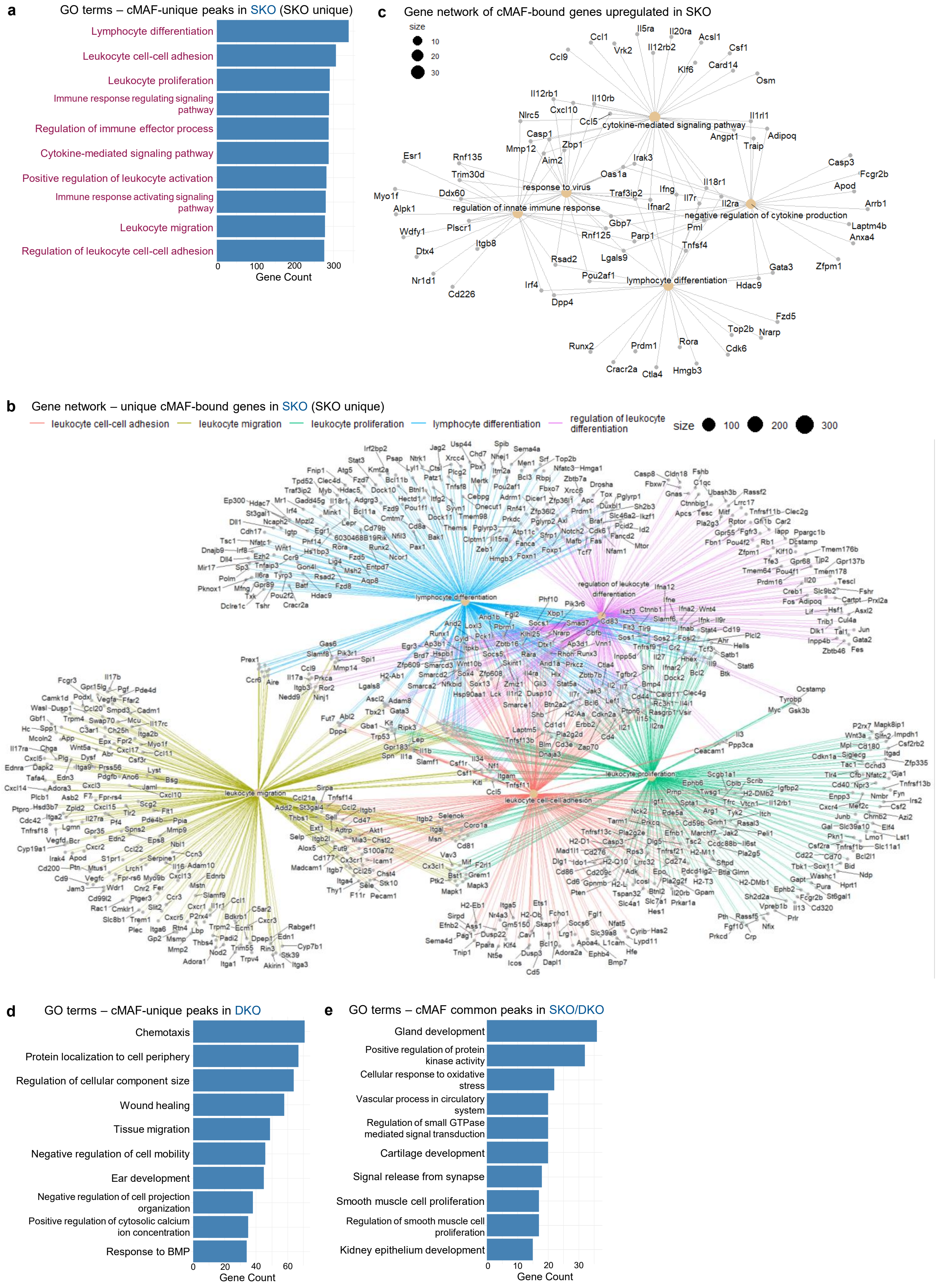

**Extended Data Fig. 7**

**cMAF chromatin occupancy in SKO and DKO thymocytes**

**a.** GO Terms of SKO-unique cMAF ChIPseq peaks in thymocytes (Hypergeometric test, FDR-adjusted pvalue, padj<0.05). **b.** Gene network of the unique cMAF-bound genes in SKO. **c.** Gene network of the cMAF-bound upregulated genes in SKO. **d.** GO Terms of cMAF-unique peaks in DKO thymocytes (Hypergeometric test, FDR-adjusted pvalue, padj<0.05). **e.** GO Terms of common cMAF ChIP-seq peaks between SKO and DKO thymocytes (Hypergeometric test, FDR-adjusted pvalue, padj<0.05)

### Extended Data Fig. 8

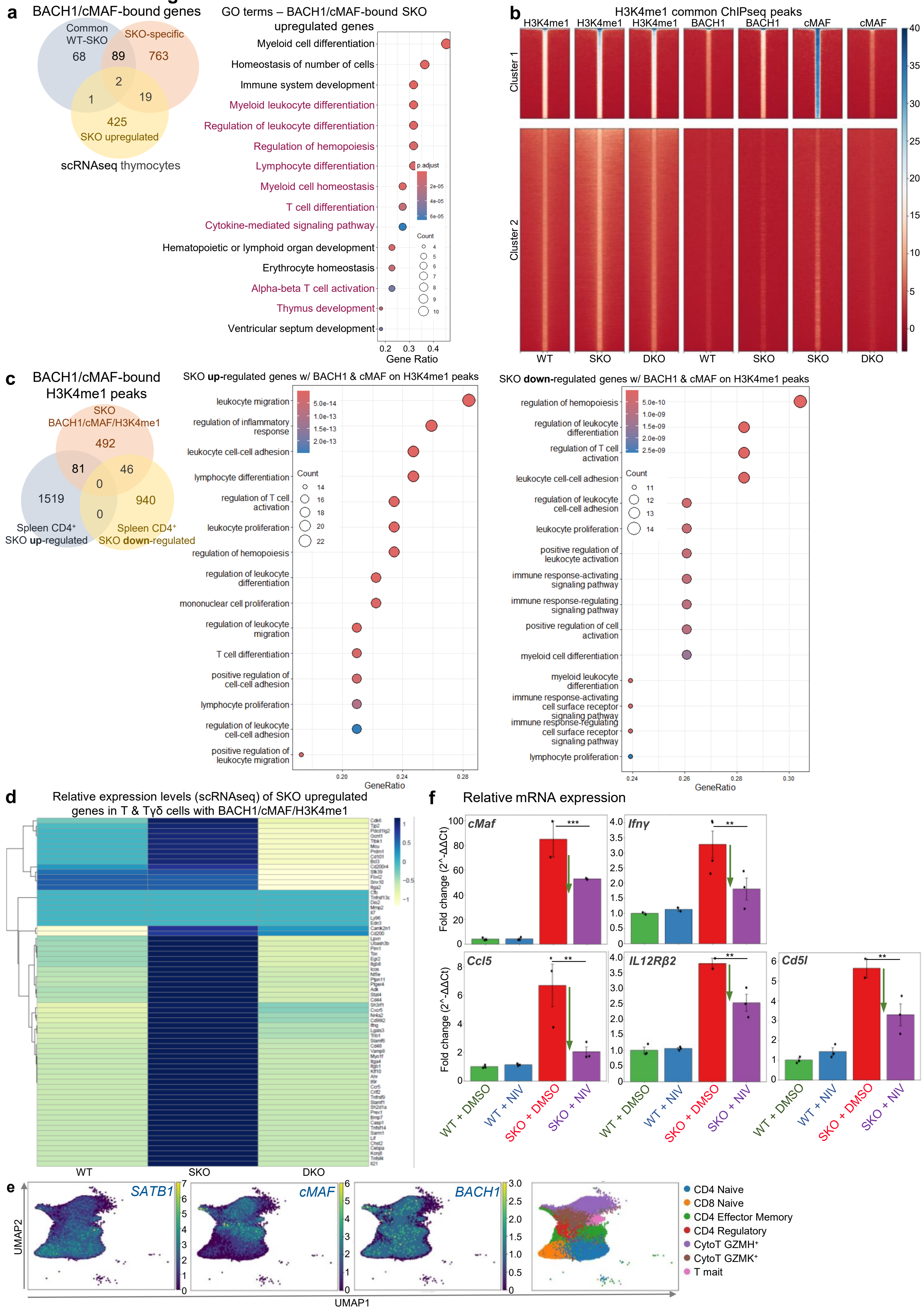

**Extended Data Fig. 8**

**Cooperative BACH1/cMAF activity drives an SLE-like gene expression program**

**a.** Venn diagram and gene ontology of the SKO upregulated genes (thymus scRNAseq) bound by BACH1 and cMAF in SKO/WT thymocytes (Hypergeometric test, FDR-adjusted pvalue, padj<0.05). **b.** K-mean clustering and heatmap of H3K4me1, BACH1 and cMAF ChIPseq binding score in the WT, SKO and DKO H3K4me1 common peaks (summit 1Kb +-5Kb). **c.** Venn diagram and gene ontology of the differentially expressed genes in WT/SKO peripheral CD4+ T cells (bulk RNAseq) that are bound by BACH1 and cMAF in H3K4me1 regions of SKO thymocytes (Hypergeometric test, FDR-adjusted pvalue, padj<0.05). **d.** Heatmap of the relative expression levels of the SKO upregulated genes (spleen scRNAseq) in WT, SKO and DKO peripheral T cells that are bound by BACH1/cMAF/H3K4me1 in SKO thymocytes. **e.** UMAP visualization of the *SATB1*, *cMAF* and *BACH1* gene expression in T cells of healthy individuals and SLE patients from scRNAseq data of PBMCs. **f.** RT-qPCR for *cMaf*, *Ifnγ*, *Ccl5*, *Il12rb2* and *Cd5l* in DMSO or Nivalenol-treated thymocytes from WT or SKO mice normalized to Hprt1 levels (ANOVA, Tukey HSD post-hoc tests).
